# A lysyl oxidase (LOX)/bone morphogenetic protein-1 (BMP1) complex to facilitate collagen remodeling

**DOI:** 10.64898/2026.03.27.714679

**Authors:** Marta Navarro-Gutiérrez, Verónica Romero-Albillo, Sergio Rivas-Muñoz, Tamara Rosell-García, Raquel Jiménez-Sánchez, Deen Matthew, Laura Marie Poller, Fernando Rodríguez-Pascual

## Abstract

Collagen biosynthesis within the extracellular matrix (ECM) relies on finely regulated enzymatic steps to ensure proper collagen maturation and fibrillar assembly. Among these, bone morphogenetic protein-1 (BMP1) and the canonical lysyl oxidase (LOX) act on the collagen telopeptide to promote procollagen processing and oxidative cross-linking, respectively. However, the mechanisms that ensure precise coordination of their activities remain poorly understood. Using NanoBiT assays, we identified and characterized a stable LOX/BMP1 protein complex that assembles intracellularly during trafficking through the ER/Golgi pathway and persists after secretion. Analysis of BMP1 and LOX domains involved in the interaction showed that BMP1 binding requires its CUB2/3 domains, while LOX recognition depends on a conserved, positively charged segment of LOX (residues 259–285) located immediately upstream of its catalytic domain. Formation of the LOX/BMP1 complex did not substantially alter LOX enzymatic activity but markedly enhanced LOX association with collagen type I through the carboxy-telopeptide region, facilitating the assembly of a ternary LOX/BMP1/collagen complex. This pre-assembled complex promoted efficient targeting of LOX to nascent collagen fibrils. Our findings reveal a previously unrecognized layer of regulation in collagen biosynthesis, in which LOX and BMP1 act as a functional unit to ensure precise localization and proper processing of collagen. This mechanism offers new insights into ECM assembly and identifies the LOX/BMP1 interface as a potentially druggable node for anti-fibrotic strategies.

## Introduction

The extracellular matrix (ECM) is a complex, three-tridimensional network that surrounds and supports cells and tissues. Beyond its structural role, the ECM is recognized as a highly dynamic system that modulates a broad range of cellular processes, including proliferation, adhesion, migration, and differentiation, and plays essential roles in tissue and organ homeostasis and regeneration. (1). The ECM is mainly composed of large, polymeric, structural proteins, such as collagens, elastin, and fibronectin, which are produced and secreted by cells in a soluble form, and subsequently assemble to form insoluble multimolecular structures. In this process, the cross-linking of molecules and fibrils by enzymatic and non-enzymatic reactions is crucial for further stabilization of these assemblies (2). In this hierarchy, lysyl oxidases (LOX) are considered the *princeps* of enzymatically mediated cross-linking effectors (3). LOX enzymes catalyze the oxidative deamination of lysine and hydroxylysine residues in collagen and elastin, yielding the corresponding aldehydes, which spontaneously condense with other oxidized groups or intact lysines to form inter- and intrachain cross-linkages. The family of LOX enzymes consists of five members in mammals (LOX, and LOX-like 1-4), with the canonical LOX being the isoform most clearly associated with the remodeling and the stability of the ECM, and described to play important roles in fibrosis, tumor microenvironment, and tissue repair (4,5).

A substantial part of our current understanding of LOX enzyme biology stems from the fact that its activity can be readily detected using *in vitro* assays with a wide variety of amine substrates, including non-peptidyl amines (6-10). However, LOX enzymatic activity in cells and tissues is specifically restricted to certain lysine (and hydroxylysine) residues of its natural substrates in the ECM. Given its apparent *in vitro* promiscuity, why doesn’t LOX indiscriminately modify amine groups in multiple proteins or metabolites *in vivo*? *In situ* activation would undoubtedly be a mechanism by which these enzymes exert a specific action on the ECM. To this respect, the canonical LOX has been reported to be proteolytically processed, and thereby activated, by bone morphogenetic protein 1 (BMP1)/Tolloid-like proteinases, a family of enzymes originally identified for their roles in C-terminal maturation of fibrillary collagen, now recognized to have numerous other substrates in the ECM (11-14). To date, however, it remains unclear where and how the protease BMP1 encounters its substrate LOX and whether this interaction effectively activates LOX enzymatic activity. Given that both proteins act on collagen telopeptides in the matrix, we hypothesized that the formation of a complex between them could promote their efficient and specific localization to sites of active matrix synthesis. In this study, we explored in greater depth the molecular dynamics of the LOX/BMP1 axis and its actions on collagen. Our results demonstrate that both proteins establish an interaction during the secretory pathway, which is preserved in the extracellular medium. We also show that the formation of this complex does not substantially alter LOX enzymatic activity, but instead facilitates its interaction with collagen *via* the telopeptide, and their subsequent incorporation into the nascent matrix. Our findings suggest the establishment of reciprocal interactions between LOX and BMP1 to promote their efficient and specific action on collagen during ECM synthesis and deposition.

## Results

### LOX and BMP1 establish a protein complex that is formed intracellularly and remains stable in the extracellular environment

To study the dynamics of LOX/BMP1, we analyzed the subcellular distribution and the potential colocalization of these proteins fused to fluorescent tags. As shown in **Figure 1A**, expression cassettes of LOX and BMP1 including their respective signal peptide sequences were C-terminally fused to green and red fluorescent proteins (GFP and RFP) to generate plasmid constructs expressing LOX-GFP and BMP1-RFP that were transiently expressed in mouse embryonic fibroblasts (MEF). **Figure 1B** shows that both proteins display a punctate pattern excluded from the nucleus that is consistent with their trafficking through the endoplasmic reticulum (ER)/Golgi system in the process of secretion to the extracellular medium. Merging both images revealed an extensive colocalization within these membranous compartments. Having demonstrated that both proteins are confined to the same spatial location, we set out to investigate whether they also interact functionally. For that purpose, we applied the NanoLuc Binary technology (NanoBiT) to reveal protein-protein interactions (PPI). This approach exploits the functional complementation of the two otherwise inactive subunits of the NanoLuc luciferase, Large-BiT (LgBiT, 18 kDa) and Small-BiT (SmBiT, 1 kDa) fused to protein partners to test their association (15). Here, we generated LOX and BMP1 versions fused at the C-terminus with LgBiT and SmBiT, respectively **(Fig. 2A)**. To account for potential non-specific interactions, we also created a control construct with a *secretable* form of the LgBiT portion by including the signal peptide sequence of the human LOX protein (sec-LgBiT, Fig. 2A). With these plasmid constructs in hand, we transfected MEF with these forms alone or with combinations of LOX-LgBiT and BMP1-SmBiT or sec-LgBiT and BMP1-SmBiT, and NanoLuc reconstitution assayed by luminometry. Expression of individual constructs yielded only residual activity slightly above the levels measured in non-transfected cells, while the combination of LOX-LgBiT and BMP1-SmBiT resulted in robust increases in NanoLuc activity, supporting the interaction between these proteins **(Fig. 2B)**. The pair sec-LgBiT/BMP1-SmBiT exhibited significantly lower NanoLuc activity, a result that reinforces the findings observed with LOX-LgBiT/BMP1-SmBiT. Similar results were observed in HEK293 cells, a cellular model previously demonstrated to faithfully recapitulate the expression and secretion of LOX and BMP1 and that has been used for further studies in this work **(Fig. 2C)** (16). As also shown in **Figure 2D**, LgBiT and SmBiT forms were efficiently expressed in HEK293 cells as detected in their expected molecular weights by immunoblotting with antibodies against NanoLuc and BMP1. Then, we aimed to confirm the interaction using an alternative approach. For that, we applied the standard co-immunoprecipitation procedure upon transfection of HEK293 cells with LOX-GFP and BMP1-RFP constructs, and captured the interaction with nanobodies against the RFP portion. As shown in **Figure 2E**, LOX-GFP was detected in the immunoprecipitate when co-transfected with BMP1-RFP, but not when transfected alone, demonstrating the interaction between these proteins.

**Figure 1.**
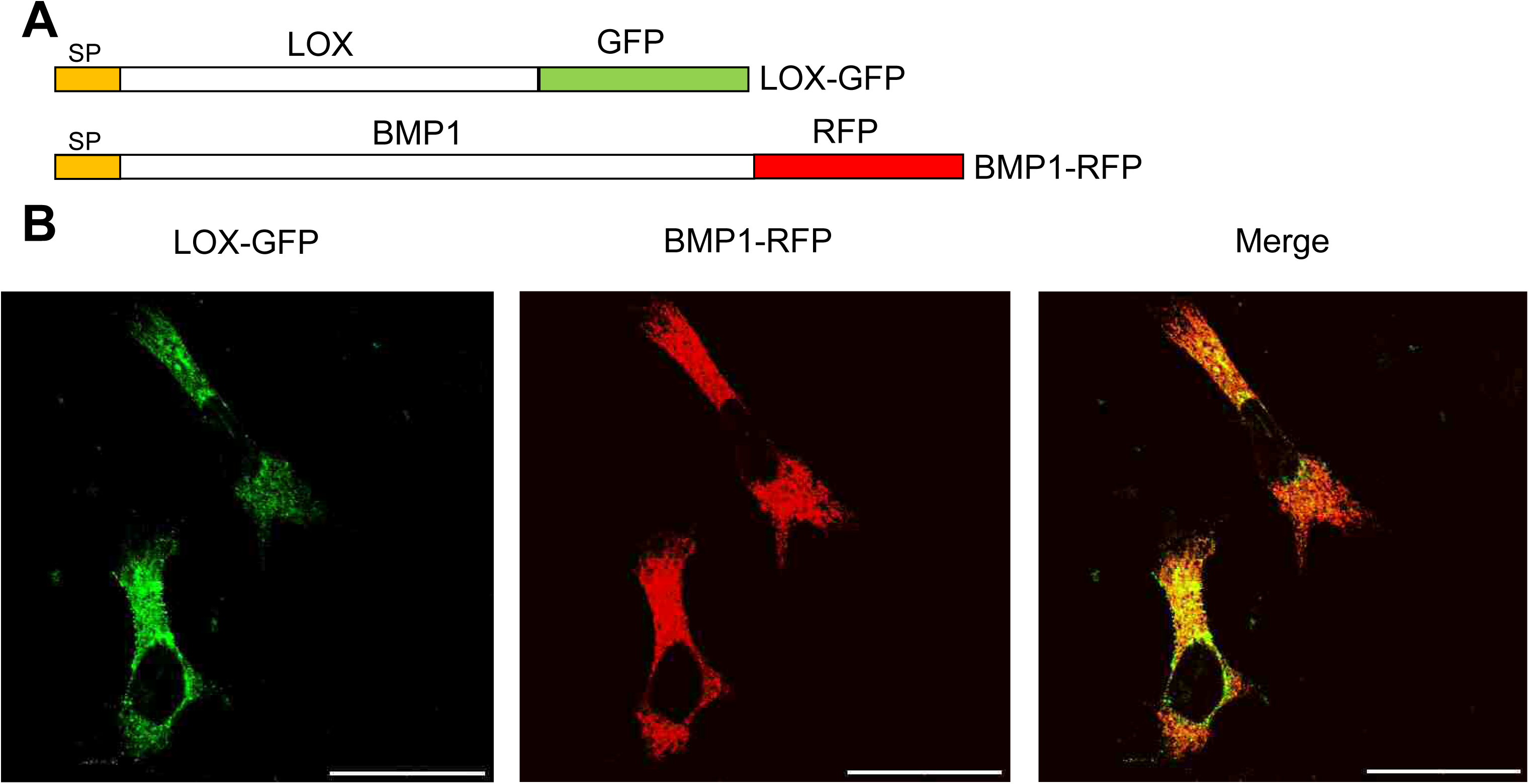
Colocalization of LOX and BMP1 in mouse embryonic fibroblasts (MEF). **(A)** Schematic representation of constructs of LOX and BMP1 fused to green and red fluorescent proteins (GFP and RFP). SP: signal peptide. **(B)** Confocal microscopy analysis of cotransfected LOX-GFP and BMP1-RFP in MEF showed extensive colocalization at the perinuclear region. Results are representative of 3 experiments. Bars=50 μm.

**Figure 2.**
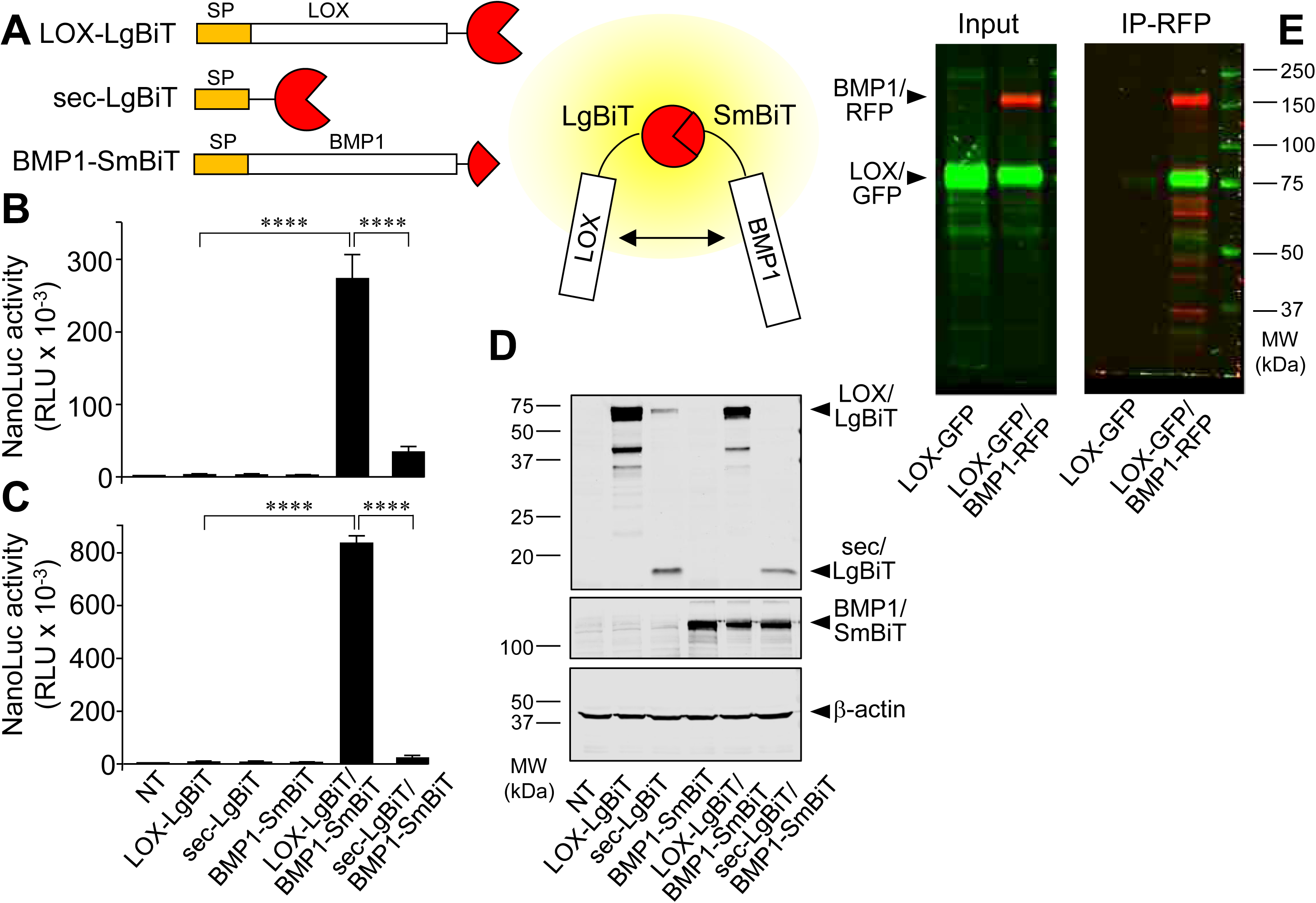
Development of a NanoBiT assay to reveal LOX-BMP1 interactions. **(A)** Schematical representation of LOX and BMP1 proteins fused at the C-terminus with Large-BiT (LgBiT) and Small-BiT (SmBiT) subunits of the NanoLuc luciferase, respectively. Interaction between these protein partners brings together LgBiT and SmBiT allowing reconstitution of the luciferase and detection of the luminescent signal. A LgBiT negative control was created that included the signal peptide (SP) sequence of the human LOX protein (sec-LgBiT). NanoLuc reconstitution assessed by luminometry in MEF **(B)** and HEK293 cells **(C)** transfected with these constructs alone or with combinations as indicated. Data are shown as relative luminescence units (RLU, mean ± s.e.m., n = 6, **** P < 0.0001; one-way ANOVA followed by Tukey’s multiple comparisons test where appropriate). NT: non-transfected **(D)** Western blot analyses of the overexpression of corresponding LgBiT- and SmBiT-tagged forms from cell lysates of HEK293 cells assayed with anti-NanoLuc (top panel), anti-BMP1 (middle panel) and anti-β-actin (bottom panel, for loading control). Arrows indicate the position of LOX-LgBiT, sec-LgBiT, BMP1-SmBiT and β-actin bands. **(E)** Standard co-immunoprecipitation analysis of HEK293 cells transfected with LOX-GFP and BMP1-RFP constructs (see Input). The immunoprecipitation (IP) panel demonstrates robust pull-down of LOX-GFP in the presence of BMP1-RFP, captured using nanobodies specific to the RFP tag. Results are representative of 3 experiments.

LOX and BMP1 are efficiently secreted to the extracellular medium (16). We were thus interested in whether the interaction is maintained in the extracellular environment. For that purpose, we transfected HEK293 cells for a cellular NanoBiT assay, but, in this case, let them express, secrete and accumulate the proteins in the extracellular medium over 48 hours. We subsequently took the cell supernatants and determined the NanoLuc activity. As can be seen in **Figure 3A**, the interaction between LOX and BMP1 remains stable in the extracellular medium. Analysis of the expressed proteins in the supernatants by immunoblotting revealed the presence of both mature and precursor forms of LOX in samples from cells transfected with LOX-LgBiT alone as a consequence of constitutive expression of BMP1 by HEK293 cells. In contrast, only the mature form was found when both proteins were co-expressed due to enhanced BMP1 expression under these experimental conditions **(Fig. 3B)**. Taken together, these results demonstrate that LOX and BMP1 form a stable complex throughout their transit along the secretory pathway and in the extracellular medium.

**Figure 3.**
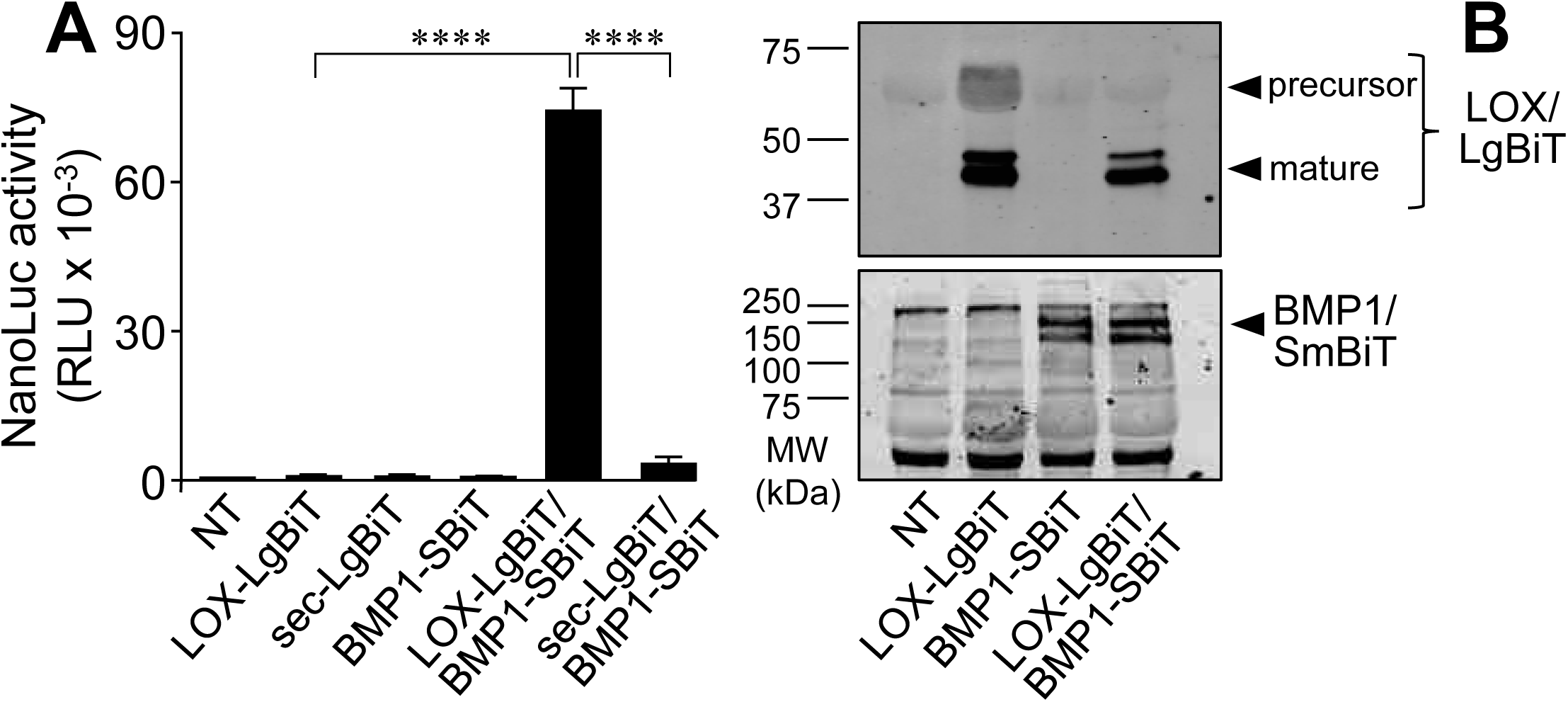
LOX and BMP1 interaction is preserved in the extracellular medium. **(A)** The NanoBiT assay was applied to cell supernatants obtained from HEK293 cells transfected with LOX-LgBiT, BMP1-SmBiT and sec-LgBiT constructs. Data are shown as relative luminescence units (RLU, mean ± s.e.m., n = 6, ****P < 0.0001; one-way ANOVA followed by Tukey’s multiple comparisons test where appropriate). **(B)** Western blots showing the expression of LOX-LgBiT and BMP1-SmBiT in cell supernatants as assayed with antibodies against NanoLuc luciferase (top panel) and BMP1 (bottom panel). Note that two forms of LOX-LgBiT, the precursor form of 65 kDa and a mature form of 47 kDa, coexist in cell supernatants from cells transfected only with LOX-LgBiT, whereas only the mature form, product of the proteolysis by BMP1, is observed in supernatants from cells co-transfected with LOX-LgBiT and BMP1-SmBiT, due to BMP1 overexpression. Results are representative of 3 experiments.

### Identification of protein domains involved in the interaction between LOX and BMP1

Having demonstrated that LOX and BMP1 form a protein complex, we aimed to identify the molecular determinants for the interaction. The mature form of BMP1 consists of an astacin-like catalytic domain followed by several CUB (for complement <u>C</u>1r/C1s, <u>u</u>EGF, <u>B</u>MP-1) and EGF (epidermal growth factor) domains (13,17). CUB domains are present in a wide variety of (mostly) extracellular and plasma membrane-associated proteins. While the general roles of CUB domains are not yet fully understood, several have been reported to be involved in oligomerization and/or recognition of substrates and binding partners (18). Considering this, we generated deletion mutants of the various CUB domains of the BMP1 protein and tested their capacity to bind LOX. Human BMP1 is expressed in two forms with variable numbers of CUB and EGF domains, which turned out to be splice variants encoded by the same gene, a long variant, known as the mammalian tolloid (mTLD) form, and a short variant, commonly referred to as BMP1 (19). Experiments shown in the previous figures of this work (Figures 1-3) were carried out with plasmid constructs corresponding to the long variant mTLD. Using this as a BMP1-SmBiT chimera, **Figure 4A and B** shows that deletion of CUB1, CUB4 and CUB5 domains did not significantly reduce NanoLuc activity and therefore alter the binding to LOX, as compared with the full-length construct. In contrast, a strong reduction in activity was observed upon the removal of CUB2 or CUB3 domains. Furthermore, we generated a construct expressing the short splicing isoform and assessed its interaction with LOX, along with that of deletion mutants derived from it lacking one or more CUB domains. As also shown in this figure, the loss of CUB2 or CUB3 was also critical for the binding **(Figure 4A and C)**. Immunoblot experiments showing the expression of these BMP1 constructs are included in **Supplementary Figure 1**.

**Figure 4.**
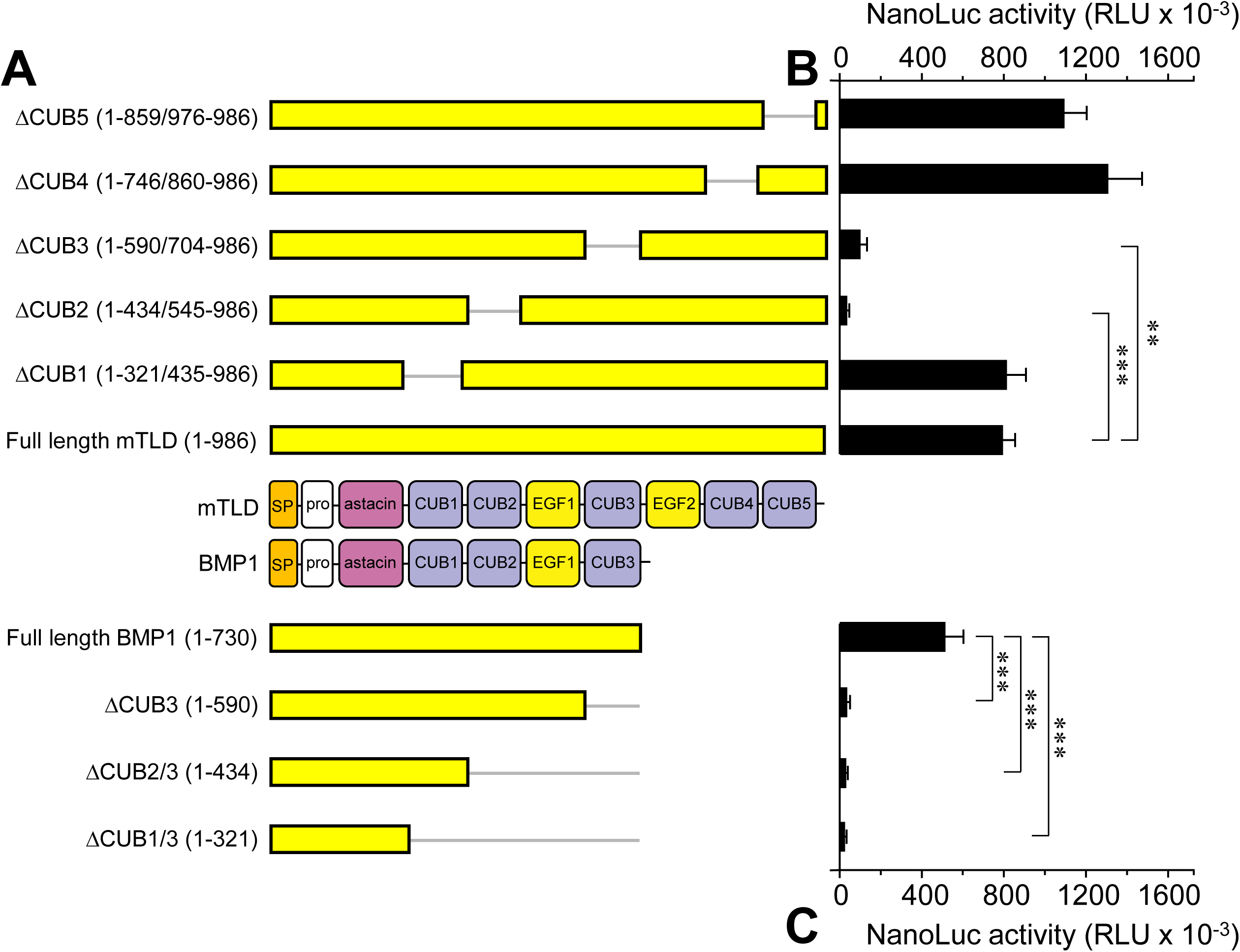
Analysis of CUB domains of BMP1 involved in the interaction with LOX. **(A)** BMP1 is expressed in two forms as a result of alternative splicing, a long variant, the mammalian tolloid (mTLD), and a short variant, commonly named as BMP1. Schematical representation of domain organization of these BMP1 forms showing the position of the astacin-like catalytic region and CUB and EGF domains. SP: signal peptide; pro: pro-peptide. SmBiT-tagged BMP1 constructs with specific deletion of CUB domains were done in the mTLD (long) and BMP1 (short) forms as indicated. NanoBiT assay from HEK293 cells transfected with mTLD **(B)** and BMP1 **(C)** constructs together with LOX-LgBiT. Data are shown as relative luminescence units (RLU, mean ± s.e.m., n = 3-6, ***P < 0.001, **P<0.01; one-way ANOVA followed by Tukey’s multiple comparisons test where appropriate).

The LOX enzyme contains two clearly distinct regions, a C-terminal portion, which includes the catalytic domain, and an N-terminal, called pro-region or pro-peptide, which is removed by BMP1 proteolysis. Homology modeling with the structure of LOXL2 in a zinc-bound precursor state, and, more recently, artificial intelligence (AI)-based AlphaFold prediction have modeled the structure of the catalytic domain with high accuracy (20,21). In contrast, the pro-region shows a low confidence score, likely adopting an unstructured, flexible conformation. To study the specific role of these regions in the formation of the complex, we have generated LOX-LgBiT variants with specific deletions of the catalytic or pro-region segments, as indicated in **Figure 5A**. NanoBiT experiments with these constructs showed that the deletion of the pro-region did not reduce the capacity of the protein to bind BMP1, whereas the loss of the catalytic domain virtually abolished the formation of the complex (**Fig. 5B and Supplementary Figure 2** for expression controls). For a more precise mapping of the sequence involved in binding, LOX-LgBiT mutants were generated that contain progressive deletions in the catalytic domain towards the N-terminus. As also shown in **Figure 5B**, variants that lost the cytokine receptor-like domain, the lysine tyrosylquinone (LTQ) cofactor anchoring or the copper-binding site, all of which are distinctive elements of the catalytic domain, retain their ability to interact with BMP1. Only when the deletion reached a 27-amino acid-long segment immediately upstream of the copper-binding site, the interaction was drastically impaired. Notably, this conserved LOX segment (residues 259–285) contains an unusually high density of positively charged residues including five arginines and one lysine, which may facilitate its interaction with the BMP1 CUB domains. These domains are known to engage in electrostatic interactions with positively charged side chains through conserved glutamate and aspartate residues in the Ca²⁺-binding site (22,23).

**Figure 5.**
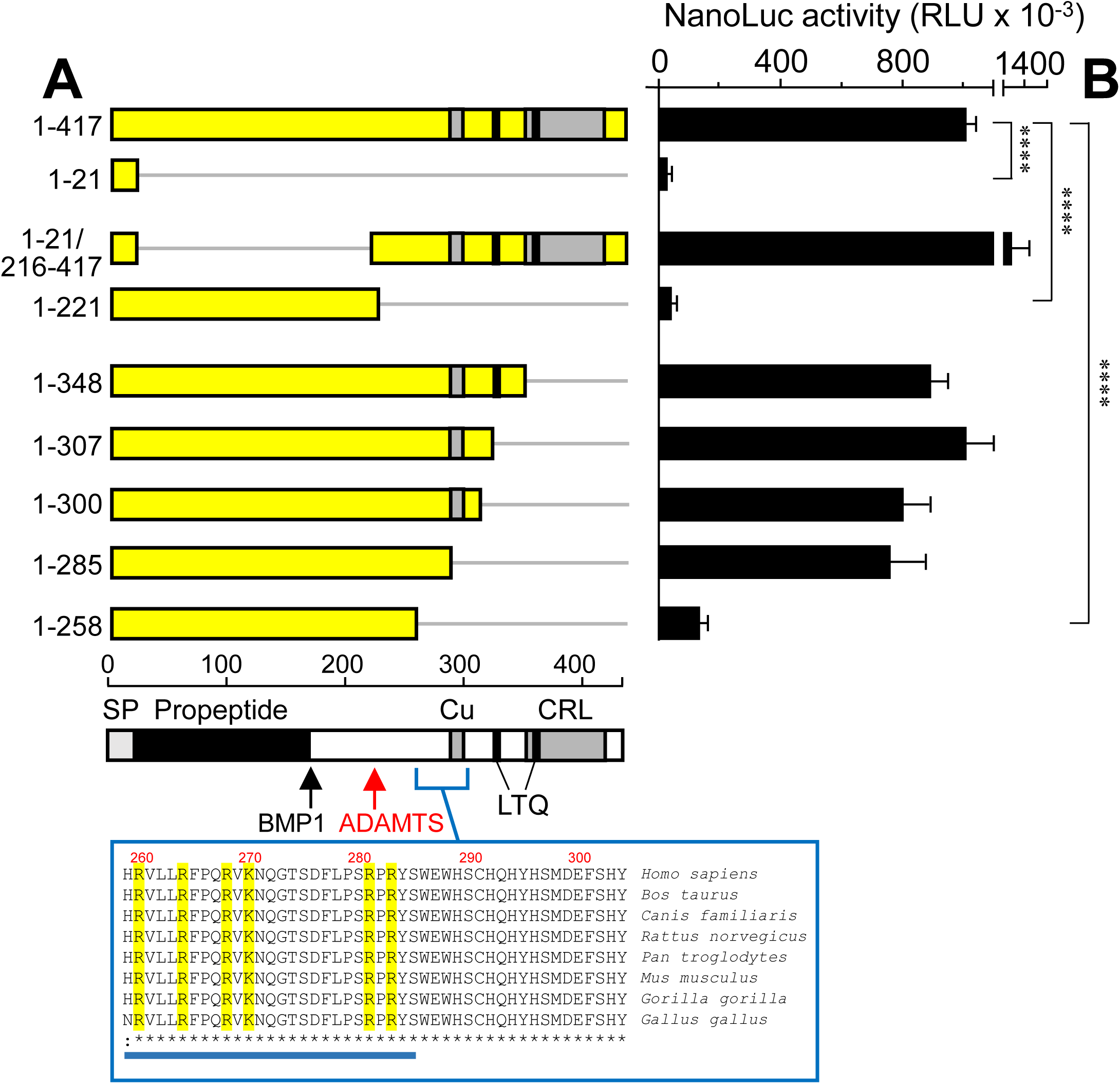
Analysis of LOX regions involved in the interaction with BMP1. **(A)** Schematical representation of the human LOX sequence highlighting the positions of the main protein regions, including the signal peptide (SP), the propeptide (black), and the catalytic domain (white). Within the catalytic domain, the cytokine receptor–like (CRL) region, the anchoring site for the lysine tyrosylquinone (LTQ) cofactor, and the copper-binding site (Cu) are shown. The proteolytic cleavage sites recognized by BMP1 and ADAMTS2/14 are also indicated. In addition, the scheme includes the LOX–LgBiT constructs containing specific deletions that were generated to delineate the protein regions involved in the interaction with BMP1. The inset shows the sequence alignment of LOX orthologs from the indicated species, illustrating the high degree of homology within the segment identified as contributing to BMP1 binding (blue bar), with the basic residues highlighted in yellow. NanoBiT assay performed in HEK293 cells transfected with the deletion **(B)** LOX-LgBiT constructs together with BMP1-SmBiT. Data are shown as relative luminescence units (RLU, mean ± s.e.m., n = 3-15, ****P < 0.001; one-way ANOVA followed by Tukey’s multiple comparisons test where appropriate).

These experiments delineated the molecular determinants underlying the interaction between LOX and BMP1, with BMP1 being either its short or long isoform. In this regard, previous studies have reported that the short isoform of BMP1 predominates in human tissues, whereas the long isoform, mTLD, is barely detectable (19). As a complementary analysis to our work, we studied whether this applies to our cell model and if this assumption can be widely generalized. To this end, we have quantified alternative splicing of the BMP1 gene by qPCR with primers amplifying exon-exon junctions for both forms in HEK293 (see **Supplementary Figure 3**). Expression levels of the long mTLD form were found to be four times higher than those of the short BMP1 one, but still within the same range. We have also surveyed public databases for RNA-seq experiments in relevant cells for ECM/fibrosis, including fibroblasts from different organs/tissues, renal mesangial or hepatic stellate cells, and found variable, but comparable, expression levels of long and short forms in the cell types analyzed (**Supplementary Figure 3 and Table 1**). Therefore, based on these evidences, both forms of BMP1 could contribute equally to LOX complex formation, which is driven by the specific interaction between the BMP1 CUB2/3 domains and a positively charged segment immediately upstream of the LOX catalytic domain.

### Biological effect of the LOX/BMP1 interaction

We were then interested in the biological role of the interaction between LOX and BMP1. We started by investigating whether BMP1 modifies the enzymatic activity of LOX. To avoid interference from endogenous expression of BMP1, we generated a BMP1 knock-out HEK293 cell clone using a CRISPR/Cas9-mediated genome editing approach, which targets exon 3 at the beginning of the astacin-like catalytic domain **(Supplementary Figure 4)**. Using this clone, we next generated a subclone expressing LOX in a tetracycline-dependent manner (referred to as KO), and compared its enzymatic activity with that of an equivalent LOX-overexpressing clone from parental HEK293 cells (referred to as WT), generated in a previous work (16). **Figure 6A** shows the expression of LOX in cell lysates and supernatants from cells incubated in the absence and presence of the tetracycline analog, doxycycline (Dox). As can be seen, both WT and CRISPR-silenced cells (KO) expressed LOX in lysates as the precursor form of 50 kDa. In the supernatants, the LOX precursor was also the predominant form, with mature bands of approximately 30 kDa only observed in WT but not in KO cells, a result consistent with the absence of BMP1 expression in these cells. We subsequently monitored enzymatic LOX activity in supernatants from Dox-induced WT and KO cells using a fluorogenic sensor (10,24). Similar activity levels were observed in both cell types, suggesting that the presence or absence of BMP1 does not significantly affect LOX activity toward the sensor **(Fig. 6B)**. LOX-dependent oxidation of the fluorescent sensor was confirmed by incubation with the pan-LOX inhibitor, β-aminopropionitrile (BAPN).

**Figure 6.**
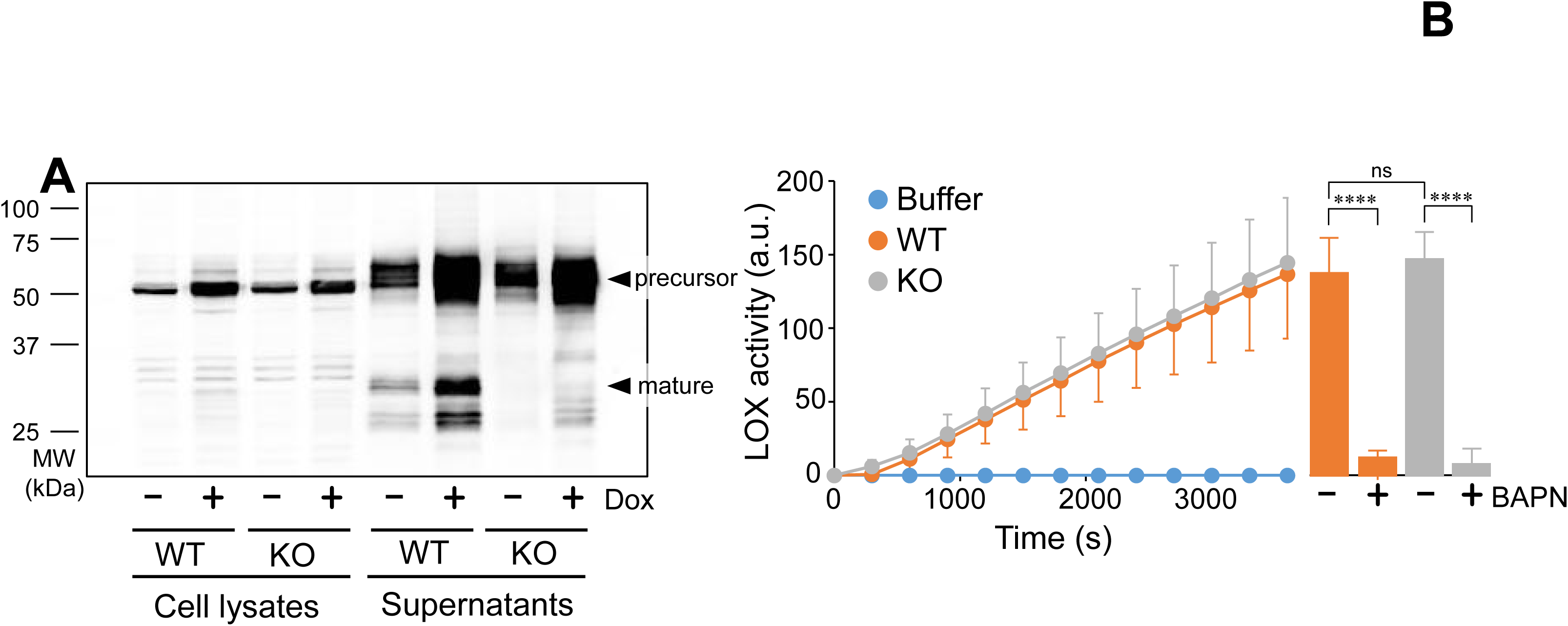
Effect of BMP1 on LOX enzymatic activity. **(A)** LOX protein species in cell lysates and supernatants from parental (WT) and CRISPR/Cas9-mediated BMP1 deletion (KO) HEK293 cells overexpressing LOX in a tetracycline-inducible manner under control conditions and upon induction with the tetracyclin analog, doxycycline (Dox), as assayed by western blot using a specific LOX antibody recognizing the C-terminal catalytic domain. Note the presence of mature forms of LOX of approximately 30 kDa in the WT but not in the KO supernatant as a result of endogenous BMP1 activity. Results are representative of 3 experiments. **(B)** Supernatants from Dox-treated WT and KO cells (or buffer PBS) were assayed with a LOX activity fluorogenic sensor in an 1-hour time-lapse assay. Graph represents arbitrary units (a.u.) of fluorescence (mean ± s.d., n=2). Bar chart corresponds to samples incubated in the presence or absence of the pan-LOX inhibitor BAPN (300 μM) and assayed with the LOX probe for one hour (a.u., mean ± s.e.m., n = 3, ****P < 0.001, ns: non-significant; one-way ANOVA followed by Tukey’s multiple comparisons test where appropriate).

Considering that LOX enzymatic activity is not substantially altered by BMP1, despite an evident proteolysis, we examined whether the formation of the LOX/BMP1 complex facilitates its interaction with collagen, a substrate of both enzymes. To address this, we set up a NanoBiT experiment that detects the interaction between LOX and collagen type I. Since both LOX and BMP1 exert their catalytic actions on the telopeptide, we expressed hemagglutinin (HA)-tagged α1 and Flag-tagged α2 forms of collagen type I containing only the C-terminal telopeptide and propeptide (Telo-Pro) segments, targeted for secretion with preprotrypsin signal sequence **(Fig. 7 and Supplementary Fig. 5A)** (25). To examine binding to the LOX-LgBiT construct, a SmBiT sequence was added to the collagen α2 chain, and experiments were done upon co-transfection with full-length BMP1 or BMP1-ΔCUB2 plasmids. Formation of the collagen type I assemblies in WT and KO cells was assessed by immunoblotting. As shown in **Supplementary Figure 5B**, HA- and Flag-immunoreactive bands of 30-35 kDa were observed in supernatants from cells transfected with Telo-Pro constructs corresponding to α1 and α2 collagen type I chains, respectively. Non-reducing electrophoresis preserved disulfide bonds and showed the formation of trimeric collagen assemblies with an apparent molecular mobility of approximately 150 kDa, which were sensitive to *in vitro* digestion with BMP1 **(Supplementary Figure 5C and D)**. After confirming the expression of this mini-collagen type I form, we assessed the formation of a putative ternary LOX/BMP1/collagen complex by NanoBiT technology both in cells and supernatants. As shown in **Figure 7B and C**, co-transfection of LOX-LgBiT with Telo-Pro in the presence of full-length BMP1 gave an increased NanoLuc signal over the background level of LOX-LgBiT alone, which was significantly reduced for BMP1-ΔCUB2, both in cells and supernatants from WT and KO HEK293 cells. Experiments performed in MEF showed essentially the same results **(Figure 7D)**.

**Figure 7.**
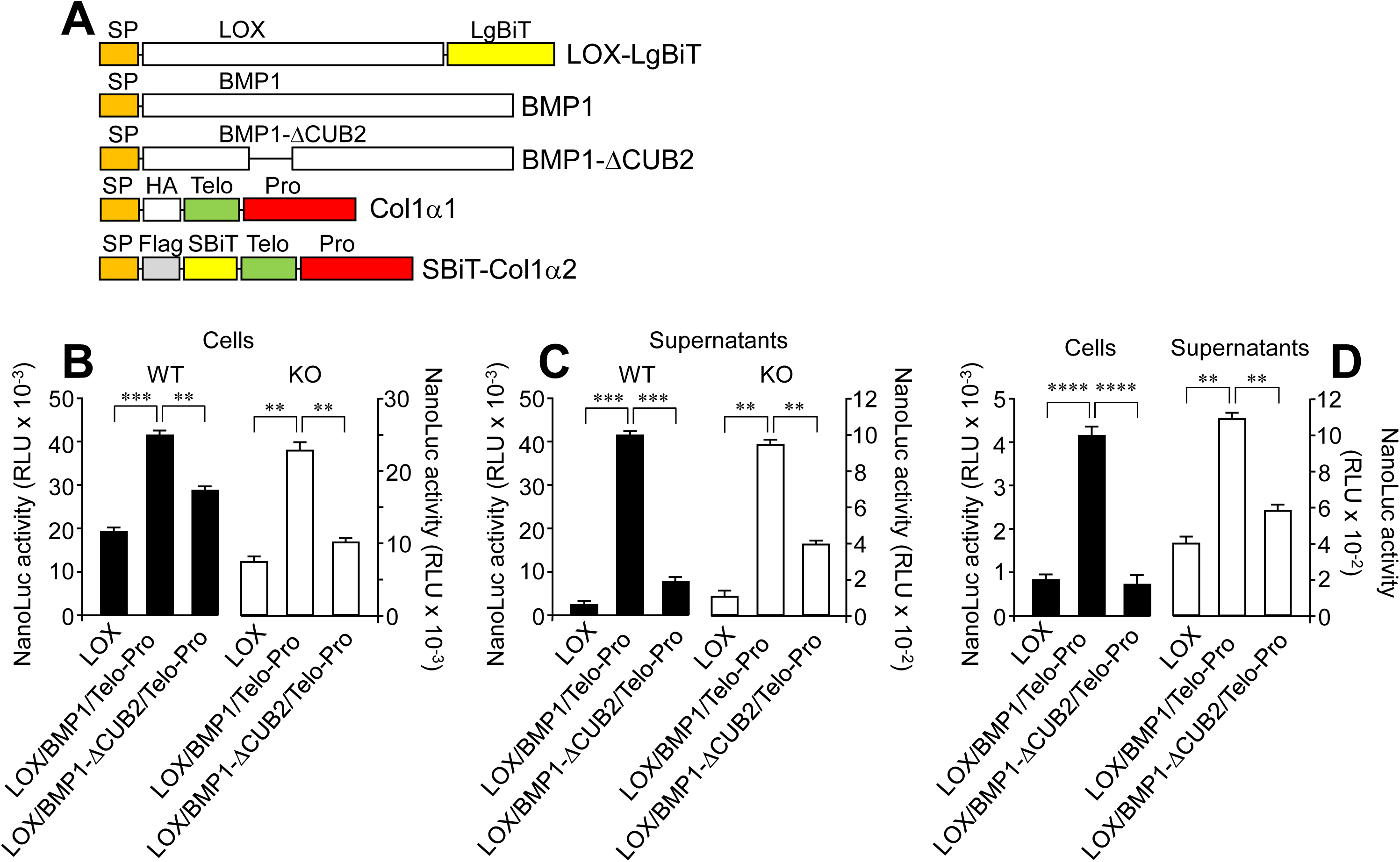
BMP1 is required for LOX to interact with collagen type I. **(A)** A NanoBiT assay was established to analyze the interaction of LOX with collagen type I using the following constructs: a LOX-LgBiT and mini-collagen construct designed for the expression of a collagen type I trimer containing only the carboxy-telopeptide and propeptide (Telo-Pro) segments, with a SmBiT sequence added in the Col1α2 form. To analyze the effect of BMP1, NanoBiT assays also included untagged full length and ΔCUB2 versions of BMP1. NanoBiT assay performed in cells **(B)** and supernatants **(C)** from WT (black bars) and KO (white bars) HEK293 cells and MEF (cells and supernatants, **D**) transfected with these constructs as indicated. Data are shown as relative luminescence units (RLU, mean ± s.e.m., n = 2-3, ****P < 0.0001, ***P<0.001, **P<0.01; one-way ANOVA followed by Tukey’s multiple comparisons test where appropriate).

Next, we tested the ability of this ternary complex to facilitate its interaction and incorporation into the ECM. For these experiments, we generated HEK293 cells stably expressing LOX-GFP in the BMP1 knockout background (LOX-GFP, KO BMP1), as well as a cell line simultaneously overexpressing LOX-GFP and BMP1-RFP (LOX-GFP/BMP1-RFP) **(Figure 8A)**. On top of the expression of these proteins, these cell clones were transiently transfected with Telo-Pro constructs. Then, the supernatants of these systems were incubated with decellularized matrix (dECM) deposited by MEF in the presence of the profibrotic cytokine transforming growth factor-β1 (TGF-β1) and with L-ascorbic acid 2-phosphate and dextran sulfate over four days **(Figure 8B)** (26). After incubation for 1 h at room temperature, unbound material was extensively washed away and retained proteins were solubilized with SDS loading buffer, boiled, and subjected to SDS-PAGE/immunoblotting to detect LOX as previously described (16). As shown in **Figure 8C**, significantly much more LOX was detected from cells capable of assembling the LOX/BMP1/collagen complex than from cells expressing only LOX and collagen, indicating a crucial role of BMP1 in promoting the formation of the ternary complex.

**Figure 8.**
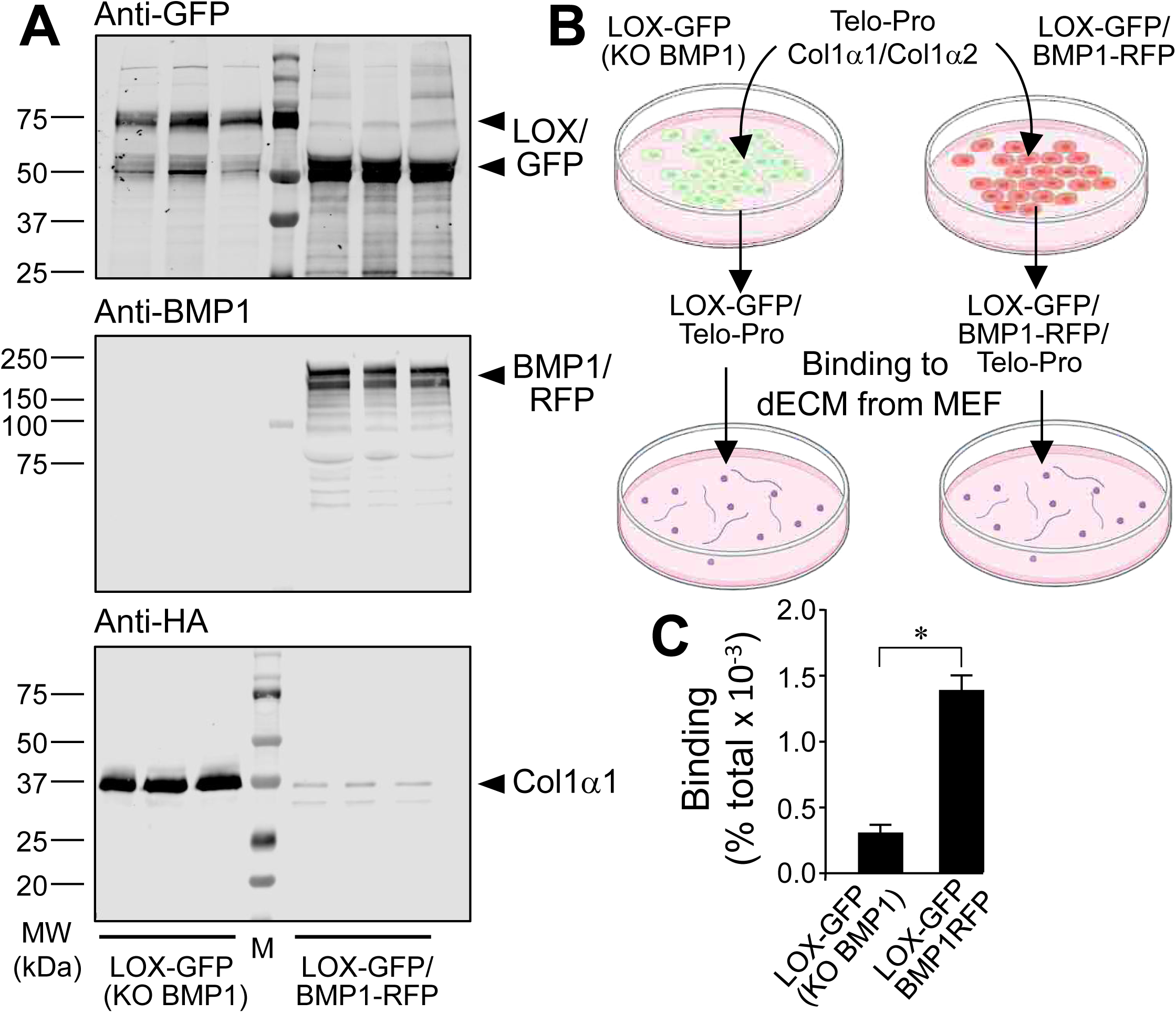
Binding assay of LOX/BMP1/collagen type I complexes to decellularized matrix. A CRISPR/Cas9 BMP1 knockout clone expressing LOX-GFP and a cell clone doubly expressing LOX-GFP and BMP1-RFP were co-transfected with Col1α1:Col1α2 Telo-Pro plasmids to generate LOX-GFP/collagen and LOX-GFP/BMP1-RFP/collagen complexes, respectively. **(A)** Characterization of the expression of LOX, BMP1 and collagen in these cells supernatants as assessed by western blot using antibodies against GFP, BMP1 and HA, as indicated. Note that the overexpression of BMP1 promotes the proteolytic processing of LOX-GFP and collagen to low molecular weight forms in LOX-GFP/BMP1-RFP, but not in LOX-GFP cells. **(B)** Schematic overview of the experiments used to analyze the capacity of protein complexes to bind decellularized matrix (dECM) from MEF. **(C)** dECM from MEF was incubated with these cell supernatants and LOX was assayed and quantified in the bound material using western blotting. Binding capacity is shown as percentage of total (mean ± s.e.m., n = 3, *P < 0.05, two-tailed unpaired t-test).

In conclusion, these experiments show that the interaction between LOX and BMP1 facilitates further formation of a complex with collagen type I via carboxy telopeptide/propeptide, that is preformed within the cells and is preserved upon secretion. The formation of this ternary complex would enable the specific targeting of LOX enzymatic activity, apparently independent of proteolysis, to collagen, likely promoting its modification and subsequent deposition more efficiently.

## Discussion

Collagen biosynthesis is a highly regulated process that requires the coordination of multiple temporally and spatially coordinated biochemical events aimed to prevent aggregation within the cell or premature association of procollagen chains in unwanted extracellular locations (27). Critical steps in its biosynthetic pathway occur both intracellularly, in the ER/Golgi, and extracellularly, where the collagen molecule undergoes several post-translational modifications, many of which are specific to collagen. As collagen folds into its triple helical structure in the ER/Golgi system, specific proline and lysine residues are modified by prolyl and lysyl hydroxylases, as well as glycosyltransferases, assisted by specialized collagen chaperones (28). After secretion into the extracellular medium, the N- and C-propeptides of procollagen are then cleaved off by a group of metalloproteinases belonging to the ADAMTS (a disintegrin and metalloproteinase with thrombospondin motifs) and BMP1 families, respectively, releasing mature collagen molecules and allowing their association into fibrils. Collagen molecules within these fibrils are then stabilized by covalent cross-linking catalyzed by LOX enzymes (5). In biological systems, enzymes catalyzing consecutive reactions within a metabolic pathway frequently assemble into multi-enzyme complexes, thereby facilitating substrate channeling and optimizing both catalytic efficiency and regulatory control of the overall process. Paradigmatic examples include the pyruvate dehydrogenase or the tryptophan synthase complexes (29-31). The collagen biosynthetic ER/Golgi machinery likewise employs this strategy. A multifunctional complex formed by prolyl 3-hydroxylase 1 (P3H1), cartilage-associated protein (CRTAP) and the cis-trans isomerase, cyclophilin B (CyPB), has been found to mediated specific prolyl-3-hydroxylation and peptidyl-prolyl cis-trans isomerization, with CRTAP playing a stabilizing role (32,33). Structural characterization of this complex has demonstrated a highly coordinated collagen flux that enables efficient prolyl residue cis–trans isomerization, hydroxylation, and molecular chaperoning, thereby ensuring proper collagen maturation (34). Here, we show that this principle extends to the final steps of collagen biosynthesis through association of LOX and BMP1 activities. Our data indicate that assembling these proteins into a single complex enhances their engagement with collagen, potentially increasing both the efficiency and specificity of their catalytic functions. We therefore propose that this complex plays a central mechanistic role in regulating collagen biosynthesis and incorporation into the nascent extracellular matrix. While LOX and BMP1 have traditionally been assumed to act exclusively in the extracellular milieu, our data further show that the complex is pre-assembled intracellularly prior to secretion, extending previous suggestions of reciprocal interactions among these ECM components during intracellular trafficking **(Proposed model in Figure 9)** (35-37).

**Figure 9.**
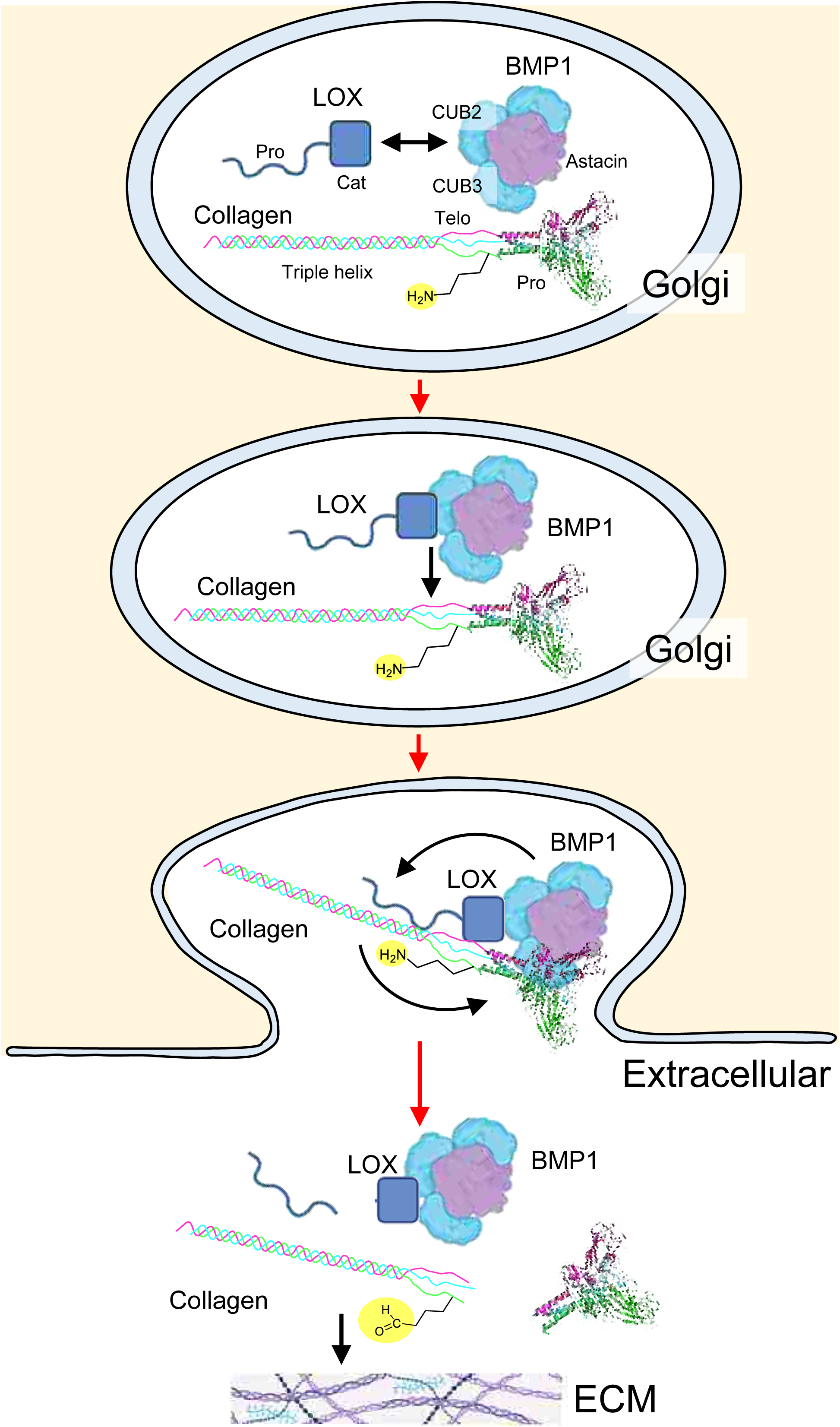
Proposed model of the interaction between LOX and BMP1 and its role in facilitating collagen remodeling. Our results suggest that LOX and BMP1 pre-assemble into a complex within the Golgi apparatus through a specific interaction between a region of LOX close to the catalytic domain and the BMP1 CUB2/3 modules. This pre-assembly enhances their subsequent engagement with collagen. Upon secretion, the resulting tripartite LOX/BMP1/collagen complex allows collagen molecules to pass through and be processed by the catalytic centers of the two enzymes, leading to C-propeptide removal and lysine oxidation within the telopeptides. These modifications render collagen competent for fibril formation and incorporation into the nascent ECM. During this cycle, LOX itself undergoes proteolytic processing by BMP1 to remove its N-terminal propeptide. For simplicity, collagen shows a representation of the triple helix and the globular C-terminal end with only one α1 telopeptide undergoing lysine oxidation.

In this study, we established a NanoBiT system to analyze the LOX/BMP1/collagen interaction network in live cells. This approach enables dynamic interrogation of the behavior of these proteins and precise dissection of the contributions of individual domains to complex formation. These advantages, however, come with limitations: the technology does not permit quantitative assessment of binding affinity or stoichiometry, which can be addressed *in vitro* using recombinant proteins. Balancing a more physiological, cell-based readout against more quantitative, protein-based assays is particularly delicate in this context. In practice, *in vitro* analyses are challenging due to the complexity of collagen, its intricate structure and extensive post-translational modifications, and the notorious difficulty of expressing and purifying LOX enzymes in standard systems. We therefore employed NanoBiT to obtain a dynamic and physiologically relevant readout of interactions within the native cellular environment, allowing us to infer mechanistic insights into collagen synthesis and deposition.

The expression and activity of LOX are intrinsically linked to severe human diseases. On the one hand, as observed in mouse KO models, deficient LOX expression or activity is associated with an altered ECM, most notably in the cardiovascular system, where it leads to severe thoracic aortic aneurysms and dissections (TAAD) (38). In this regard, natural LOX variants identified as causative in TAAD disrupt the catalytic domain, either through frameshift/nonsense mutations or by missense substitutions affecting residues essential for catalysis (39-43). Notably, some pathogenic variants map directly to the LOX protein segment characterized in this study (41). These variants could specifically disrupt the LOX/BMP1 interaction and, consequently, impair collagen maturation. On the other hand, conditions marked by increased collagen synthesis and deposition, accompanied by elevated LOX expression and/or activity, arise in fibrosis, a pathological feature of many chronic inflammatory diseases associated with substantial global morbidity and mortality (44). Although significant progress has been made in elucidating the mechanisms underlying fibrosis, there remains a critical shortage of clinically translatable compounds with robust anti-fibrotic activity (45). Thus, despite new additions to the therapeutic portfolio, current pharmacological options primarily target signaling effectors downstream of pro-fibrotic pathways, achieving only modest efficacy and with indications largely limited to idiopathic pulmonary fibrosis, the most prevalent form (46,47). Ideally, a drug that modulates collagen production by activated fibroblasts or myofibroblasts may be highly valuable for treating fibrosis across diverse tissues and disease contexts. Based on our results, the interaction between LOX and BMP1, and the subsequent formation of a ternary complex with collagen, constitutes a novel therapeutic target for the development of specific disruptors with anti-fibrotic activity, potentially offering greater specificity than inhibitors targeting LOX or BMP1 alone.

In summary, our study reveals that LOX and BMP1 form a functional complex, mediated by the BMP1 CUB2/3 domains and a conserved, positively charged LOX segment immediately upstream of the catalytic domain, that enhances their coordinated engagement with collagen and presumably promotes efficient maturation of the ECM. Evidence of intracellular pre-assembly further reframes where and how these interactions are orchestrated, complementing the traditional view of exclusively extracellular processing. Future work integrating quantitative binding assays with *in vivo* perturbations will be essential to define stoichiometry, temporal ordering, and therapeutic tractability of this complex.

## Experimental Procedures

### Cell culture

Mouse embryonic fibroblasts (MEF) were kindly provided by Dr. Inmaculada Navarro-Lérida (Centro de Biología Molecular Severo Ochoa, Spain) and cultured in Dulbeccós modified Eagle medium (DMEM) supplemented with 10% fetal bovine serum (FBS), 100 U/ml penicillin and 100 μg/ml streptomycin at 37°C in a humidified 5% CO_2_-95% air incubator. Human embryonic kidney 293 (HEK293) and Flp-In T-REx 293 cells (Invitrogen) cells were maintained in culture according to protocols previously published (48).

### Fluorescence microscopy

To study potential colocalization of LOX and BMP1 proteins, we generated constructs for the overexpression of fusion proteins of human LOX and BMP1 with green fluorescent protein (GFP) and red fluorescent protein (RFP), respectively. For that purpose, GFP and RFP cassettes were C-terminally introduced into previously described pcDNA5/FRT/TO-LOX and pcDNA5/FRT/TO-BMP1 plasmids to obtain the corresponding pcDNA5/FRT/TO-LOX-GFP and pcDNA5/FRT/TO-BMP1-GFP **(see Figure 1A)** (16). These vectors, which contain the native signal sequence of their genes, were then co-transfected in MEF seeded on glass coverslips using Lipofectamine LTX and Plus Reagent (Invitrogen) and fusion proteins expressed for a 24-hour period before imaging. Cells were examined with a LSM900 Zeiss confocal microscope with the Zeiss Zen software for acquisition image. GFP was excited by a 488 nm (blue) laser with 505-530 emission, while RFP was excited with a 555 nm (yellow green) laser with 560-615 emission using a 40x objective. Images were processed by Fiji software image analysis.

### Analysis of protein-protein interactions by the NanoBiT technology

Assembly of protein complexes was assayed with NanoBiT (Promega), a complementation system built on NanoLuc luciferase, using methods previously described (15,49). Briefly explained, the strategy is based on the splitting of NanoLuc into two segments, a large 18 kDa fragment called Large BiT (LgBiT) and a much smaller 1 kDa (11 aminoacids) fragment named Small BiT (SmBiT). The fragments present low intrinsic affinity for each other but reconstitute the full bioluminescent protein when brought together. The system exploits these properties by fusing LgBiT and SmBiT to potential protein partners. Here we have fused both LgBiT and SmBiT portions C-terminally of LOX and BMP1 in order to detect interactions between these proteins **(see Figure 2A)**. For that, LgBiT and SmBiT sequences preceded by the flexible linker SSGGGGSGGGGSSG were inserted downstream LOX and BMP1 cassettes in pcDNA5/FRT/TO-LOX and pcDNA5/FRT/TO-BMP1 plasmids by standard cloning methods. A control vector for the expression of a *secretable* form of the LgBiT protein was generated with a LgBiT cassette including the signal peptide sequence of the human LOX protein (sec-LgBiT). To further analyze potential interactions with collagen type I, plasmids for expression of hemagglutinin (HA)-tagged C-Pro α1(I) and Flag-tagged C-Pro α2(I) forms of collagen type I with a preprotrypsin signal sequence were kindly provided by Matthew D. Shoulders (Massachusetts Institute of Technology, Cambridge, USA). These constructs were modified to include the telopeptide sequence in both plasmids and a N-terminal SmBiT portion in the α2(I) form **(see Figure 7A)**. These constructs were introduced in cells in a ratio 2:1 between α1(I) and α2(I) chains giving to the expression and secretion of a functional SmBiT-containing Telo-Pro trimer assembly. LOX-LgBiT and BMP1-SmBiT (and BMP1 alone) constructs with domain deletions were generated by PCR-based site-directed mutagenesis as previously described (48). All constructs were verified by sequencing.

MEF or HEK293 cells were transfected with these plasmids either individually or in combinations LgBiT/SmBiT using Lipofectamine 2000 (Invitrogen). NanoLuc reconstitution was monitored using the Nano-Glo® Live Cell assay kit (Promega) in cells after 24 hours or accumulated in the supernatants for 48 hours (49).

### Protein analysis

Protein expression was analyzed by SDS-polyacrylamide electrophoresis coupled to immunoblotting. For these experiments, cell monolayers were washed with phosphate buffered saline (PBS) and lysed with 750 µl Tris-SDS buffer [60 mM Tris-HCl, pH 6.8, 2% sodium dodecyl sulphate (SDS)] to obtain total cell lysate. Proteins accumulated in cell supernatants were collected and concentrated using Amicon Ultracentrifugal filters (Ultracel-10K, EDM Millipore). Proteins were fractioned on SDS-polyacrylamide gels using a Mini Protean electrophoresis cell (Bio-Rad) and then transferred onto nitrocellulose membranes at 12 V for 20 min in a semi-dry Trans-Blot Turbo system (Bio-Rad). Membranes were blocked by incubation for 30 min with 1% bovine serum albumin (BSA) in Tris-buffered saline (TBS) containing 0.5% Tween-20, and antigens were detected using specific primary antibodies (NanoLuc: MAB10026, R&D Systems; BMP1: AF1927, R&D Systems; LOX: ab31238, Abcam; GFP: Cat.No. 11814460001, Sigma; RFP: 5F8, Chromotek; hemagglutinin (HA): 16B12, Biolegend; Flag: 12CA5, Sigma; β-actin: A5441, Sigma). Blots were then developed using corresponding IRDye 680 or 800 labeled secondary antibodies with the Odyssey M infrared imaging system (LI-COR).

For immunoprecipitation experiments, HEK293 cells transfected with LOX-GFP or LOX-GFP/BMP1-RFP were washed once with ice-cold PBS and immediately lysed with RIPA buffer (50 mM Tris-HCl, pH 8.0, with 150 mM sodium chloride, 1.0% Nonidet P-40, 0.5% sodium deoxycholate, and 0.1% sodium dodecyl sulfate). Lysates were cleared from cell debris by centrifugation at 14,000xg for 20 min at 4°C and protein concentration in the supernatants was determined by the BCA assay. Immunoprecipitation studies were done by incubation of protein extracts with RFP-Trap® agarose (Chromotek) in RIPA buffer for 1 hour on ice according to manufacturer’s instructions. Captured proteins were analyzed by immunoblotting with anti-GFP and –RFP antibodies as described above.

### Analysis of alternative splicing of mTLD/BMP1

Alternative splicing of the mTLD/BMP1 gene in HEK293 cells was analyzed by quantitative real time reverse-transcribed polymerase chain reaction (qRT-PCR) as previously described (50). Briefly, total RNA from HEK293 cells was extracted using the RNAeasy kit (Qiagen) and used for cDNA synthesis with the High Capacity cDNA Reverse Transcription Kit (Applied Biosystems). qRT-PCR was performed using iQ SYBR Green Supermix (Bio-Rad) with primers amplifying specific exon-exon junctions for mTLD (long) and BMP1 (short) forms in a CFX96 thermocycler (Bio-Rad) **(Supplementary Figure 3)**. Expression levels were quantified using a standard curve generated from plasmid DNA vectors encoding the mTLD and BMP1 forms, and relative abundances were calculated accordingly.

Additionally, publicly available RNA-seq datasets generated from ECM- and fibrosis-relevant cell types, including fibroblasts from different tissues, renal mesangial cells, and hepatic stellate cells, were analyzed to quantify the relative abundance of both transcripts. For this purpose, GEO project identifiers were used to identify samples from the selected studies and to retrieve the corresponding SRA accession numbers for each sample. Raw sequencing data were then downloaded from SRA as FASTQ files using fasterq-dump from the SRA-toolkit (version 3.0.1) and data quality assessed using FastQC (version 0.11.9). Upon extracting reduced read quality samples, trimming was performed using Trim Galore (version 0.6.10) and reads aligned to the human reference genome assembly (GRCh38.p14; *Homo_sapiens*) using HISAT2 (version 2.1.0). Sashimi plots were then generated from these alignments using ggsashimi and number of reads overlapping each junction used to estimate the relative abundance of the two isoforms (51).

### CRISPR/Cas9 gene editing

CRISPR/Cas9-mediated gene deletion of the BMP1 gene in Flp-In T-REx 293 cells (Invitrogen) was performed using the protocol described by Ran *et al*. (52). Briefly, a human BMP1-targeting single-guide RNA (sgRNA) previously reported by Ramakrishna *et al.* was cloned into the vector pSpCas9(BB)-2A-Puro (Addgene #62988), which features a Cas9 expression cassette and puromycin resistance **(see Supplementary Figure 4A)** (53). Upon transfection, cells were challenged with puromycin and surviving colonies isolated and expanded for screening by immunoblotting. Genomic DNA sequence around the target site was amplified by PCR using primers previously reported, cloned into pCR2.1 vector using a TA-cloning kit (Invitrogen), transformed and grown in *E. coli* bacteria and sequenced (53). On top of this BMP1 null background, a tetracycline-inducible cell clone stably expressing LOX was generated using the Flp-In T-REx 293 system (16). Briefly, pcDNA5/FRT/TO and Flp recombinase expression plasmid pOG44 were cotransfected using Lipofectamine 2000 (Invitrogen). After 48 h, cells with insertion of LOX cDNA into FRT sites were selected with hygromycin B (Invitrogen). Isogenic pooled clones were expanded and checked for transgene expression by immunoblotting of cell lysates and supernatants after 48 hours of incubation in the presence and absence of the tetracycline analog, doxycycline, at 1 μg/ml. A similar LOX-overexpressing cell clone in parental Flp-In T-Rex 293 cells used as control was previously reported (16).

### Lysyl oxidase (LOX) enzymatic activity

LOX activity in cell culture supernatants was measured using a recently developed Pacific Blue–derived LOX activity sensor (10). In brief, concentrated supernatants or PBS buffer were incubated with the LOX sensor (0.5 mM final concentration), and fluorescence intensity (excitation at 365 nm; emission 410–460 nm) was recorded every 300 seconds for one hour in a total volume of 50 μl using a Glomax Multi Detection System (Promega). Parallel assays were performed in the presence of 300 μM β-aminopropionitrile (BAPN), a pan-LOX inhibitor, to confirm assay specificity. Enzyme activity was expressed in arbitrary fluorescence units.

### Analysis of binding to decellularized matrix

The ability of the protein complexes to bind the ECM was assessed by incubation of MEF-derived decellularized matrices generated under macromolecular crowding conditions with cell culture supernatants and analysis of the bound material using immunoblotting as previously described (16). To obtain supernatants enriched in protein complexes, HEK293 cells expressing LOX–GFP in a BMP1-null background, or co-expressing LOX–GFP and BMP1–RFP, were generated using the Flp-In T-REx 293 system. For the dual-expression cell line, a pcDNA5-FRT-TO vector containing both LOX–GFP and BMP1–RFP expression cassettes was engineered using appropriate restriction enzyme sites. Once the stable clones were established, cells were transiently transfected with Telo-Pro plasmids using Lipofectamine 2000 to promote assembly formation. MEF decellularized matrices were produced by culturing MEF in serum-free medium supplemented with 100 μg/ml 500 kDa dextran sulfate (DxS) and 29 μg/ml L-ascorbic acid 2-phosphate (Sigma-Aldrich), in the presence of 5 ng/ml TGF-β1 (R&D Systems) for four days. Decellularization was performed by treating the cultures with extraction buffer containing 0.5% (v/v) Triton X-100 and 20 mM NH₄OH in PBS for 3–5 minutes, as previously described (54). Supernatants were then incubated with the decellularized matrices for 1 hour at room temperature. Unbound proteins were removed by extensive washing with PBS, and matrix-bound material was eluted by adding 30 μl SDS loading buffer followed by 5 minutes of heating at 60 °C and assayed for LOX by immunoblotting with anti-LOX antibody. LOX levels were quantified by densitometry, with binding expressed as percentage of LOX input.

### Statistical analysis

Experimental data were analyzed using GraphPad Prism version 11.0.0 by one-way ANOVA followed by Tukey’s test for multiple comparisons, or using two-tailed unpaired t-test for comparison between two groups. Adjusted or standard P values are indicated in the figure legends where appropriate (statistically significant when P<0.05).

## Acknowledgements

This work was supported by MICIU/AEI/10.13039/501100011033 through grants PID2022-136703OB-I00 (ERDF A way of making Europe) and CPP2022-009582 (European Union NextGenerationEU/PRTR), and a predoctoral fellowship of the FPI Program (PREP2022-00714, Marta Navarro-Gutiérrez), and by Comunidad de Madrid INNOREN-CM S2022/BMD7221. The CBMSO is supported by the Severo Ochoa Centers of Excellence Program (MICIU/AEI/10.13039/501100011033) and receives institutional support from Fundación Ramón Areces. The BMP1 alternative splicing data analysis has been performed by the Biocomputational Analysis Core Facility (http://www.cbm.uam.es/genomica) at the CBMSO. The Biocomputational Analysis Core Facility also receives funding from the European Commission – NextGenerationEU, through Momentum CSIC Programme: Develop Your Digital Talent. Special thanks to María Santos Romero. Imaging studies has been performed in the Advanced Light Microscopy Facility at the CBMSO. We thank Inmaculada Navarro-Lérida (CBMSO) for providing mouse embryonic fibroblasts (MEF) and Matthew D. Shoulders (Massachusetts Institute of Technology, Cambridge, USA) for sharing plasmids for the expression of HA-tagged C-Pro α1(I) and Flag-tagged C-Pro α2(I) forms of collagen type I. We thank Javier García-Marín (University of Alcalá de Henares, Madrid) and Helma Wennemers (ETH Zurich) for his helpful comments and suggestions to improve this work.

## Author Contribution Statement

Fernando Rodriguez-Pascual and Marta Navarro-Gutierrez delineated the experimental plan, performed most of the experiments, analyzed the data and contributed to manuscript text and reviewed its final version. Verónica Romero-Albillo, Sergio Rivas-Muñoz, Tamara Rosell-García and Raquel Jiménez-Sánchez carried out different experiments. Deen Matthew and Laura Marie Poller generated the LOX activity sensor, developed its experimental application, and contributed to manuscript text and reviewed its final version.

