## Supplementary Material for "A lysyl oxidase (LOX)/bone morphogenetic protein-1 (BMP1) complex to facilitate collagen remodeling"

[1]Marta Navarro-Gutiérrez, [1]Verónica Romero-Albillo, [1]Sergio Rivas-Muñoz, [1]Tamara Rosell-García, [1]Raquel Jiménez-Sánchez, [2]Deen Matthew, [2]Laura Marie Poller and [1]Fernando Rodríguez-Pascual[*]

[1]Centro de Biología Molecular Severo Ochoa, Consejo Superior de Investigaciones Científicas (CSIC)/ Universidad Autónoma de Madrid (UAM), Madrid, Spain.
[2]Laboratory of Organic Chemistry, ETH Zürich, Switzerland.

Correspondence footnote:
Dr. Fernando Rodríguez-Pascual
Centro de Biología Molecular Severo Ochoa (CBMSO)
Consejo Superior de Investigaciones Científicas (CSIC)/Universidad Autónoma de Madrid (UAM).
Nicolás Cabrera, 1
E-28049,
Madrid, Spain


**Supplementary Figure 1. Expression control for BMP1 forms.** Western blot analysis of the expression of the mTLD **(A)** and BMP1 **(B)** forms fused to SmBiT used in Figure 4 as assayed with a specific antibody against BMP1. Blots were also probed with an anti-β-actin antibody to confirm equal loading. Results are representative of 3 experiments.

**Supplementary Figure 2. Expression control for LOX forms. (A and B)** Western blot analysis of the expression of deletion mutants of LOX fused to LgBiT used in Figure 5 as assayed with a specific antibody against NanoLuc. Blots were also probed with an anti-β-actin antibody to confirm equal loading. Results are representative of 3 experiments.

**Supplementary Figure 3. Analysis of alternative splicing of BMP1 gene. (A)** Schematical representation of splicing events in the human BMP1 gene with indications of the exon usage and the resulting mTLD and BMP1 forms, as well as the positions of primers used **(B)** for characterization in HEK293 cells. **(C)** Abundance levels of mTLD (long)/BMP1 (short) forms in different cells relevant for ECM/fibrosis as inferred from public databases of RNA-seq experiments (detailed information in Table I). Upper bar of this graph corresponds to values obtained experimentally by qPCR-RT from HEK293 cells in our work. Results are representative of 3 experiments.

**Supplementary Figure 4. CRISPR/Cas9-mediated deletion of BMP1 gene in HEK293 cells. (A)** Schematical representation of the CRISPR/Cas9 design with target in exon 3 of human BMP1 gene. **(B)** BMP1 expression in isolated CRISPR/Cas9 BMP1 deleted cell clones as compared with parental HEK293 cells (HEK293) and one clone overexpressing BMP1 (BMP1). Note that all clones except one (clone #6) were devoid of BMP1 expression. M: molecular weight marker. Clone #1 was selected for further experiments and genotyped by PCR and Sanger sequence analysis (see in A).

**Supplementary Figure 5. Expression control for mini-collagen type I. (A)** Schematical representation of mini-collagen constructs designed for the expression of a collagen type I trimer assembly containing the carboxy-telopeptide and propeptide (Telo-Pro) segments (α1: HA tag; α2: Flag and SmBiT tags). Detection of Col1α1 and Col1α2 chains (**B**: reducing; **C**: non-reducing conditions) by western blot with anti-HA and -Flag, respectively, in non-transfected (NT) and transfected (Telo-Pro) parental WT and KO (CRISPR/Cas9 BMP1 deleted) HEK293 cells. **(D)** *In vitro* proteolysis by BMP1 of mini-collagen type I as assessed by Western blot with anti-HA and -Flag antibodies. Results are representative of 3 experiments. Immunoreactive bands detected with the anti-Flag antibody at higher molecular weights than Telo-Pro Col1α2 were observed in the KO cells (both transfected and non-transfected), as they originate from the vector used for CRISPR/Cas9 editing.

**Table I. Summary of the RNA-seq datasets analyzed for alternative splicing of the BMP1 gene.** The table lists the corresponding GEO Project ID, the Sequence Read Archive (SRA) project and identifier, a description of the cell type/tissue, and the percentages of BMP1 (short) and mTLD (long) transcripts for each sample, along with the overall average.

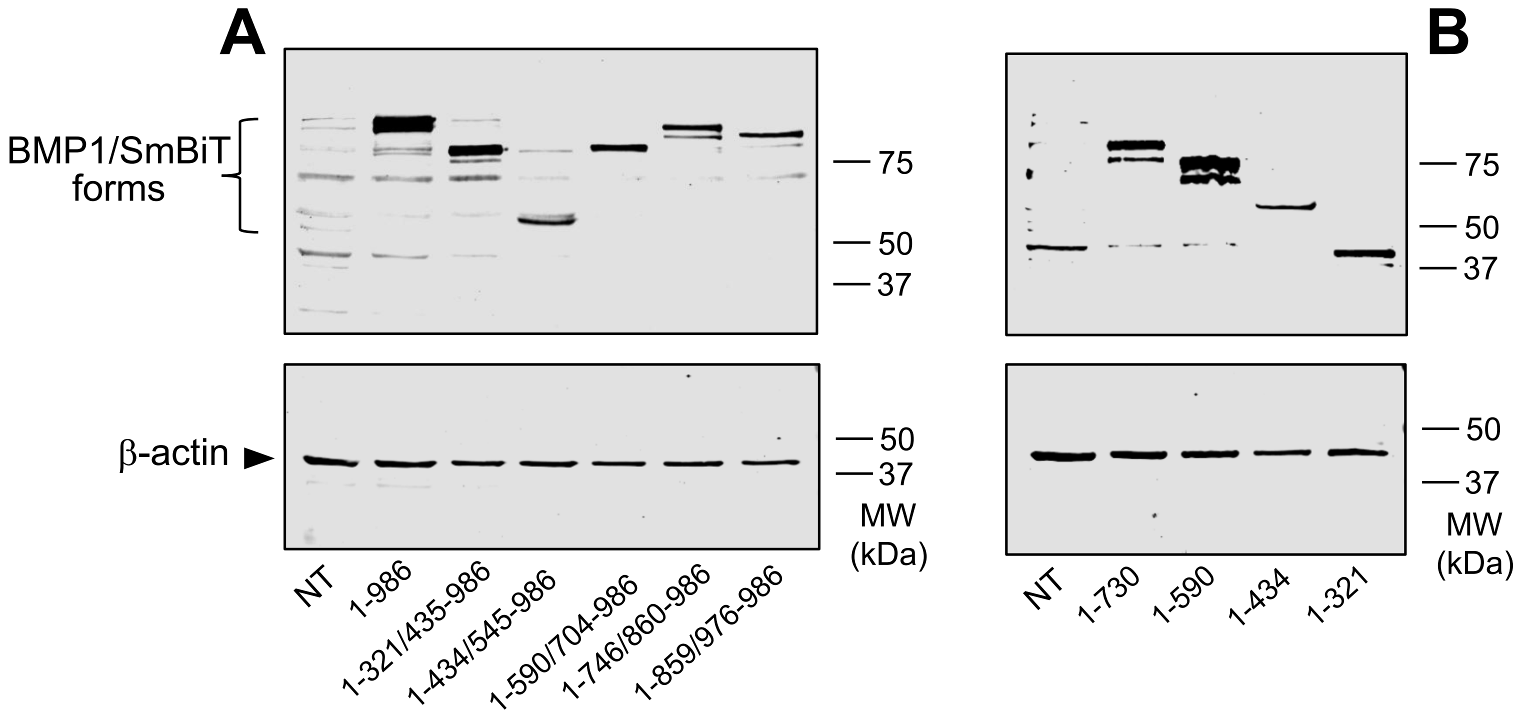


**Supplementary Figure 1**

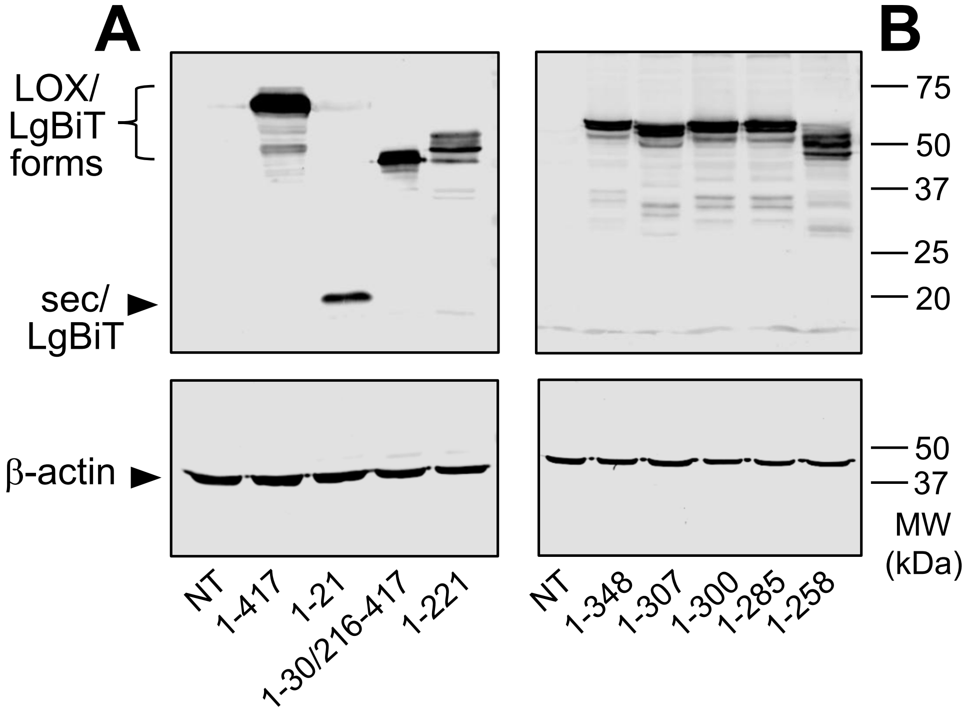


**Supplementary Figure 2**

**A**

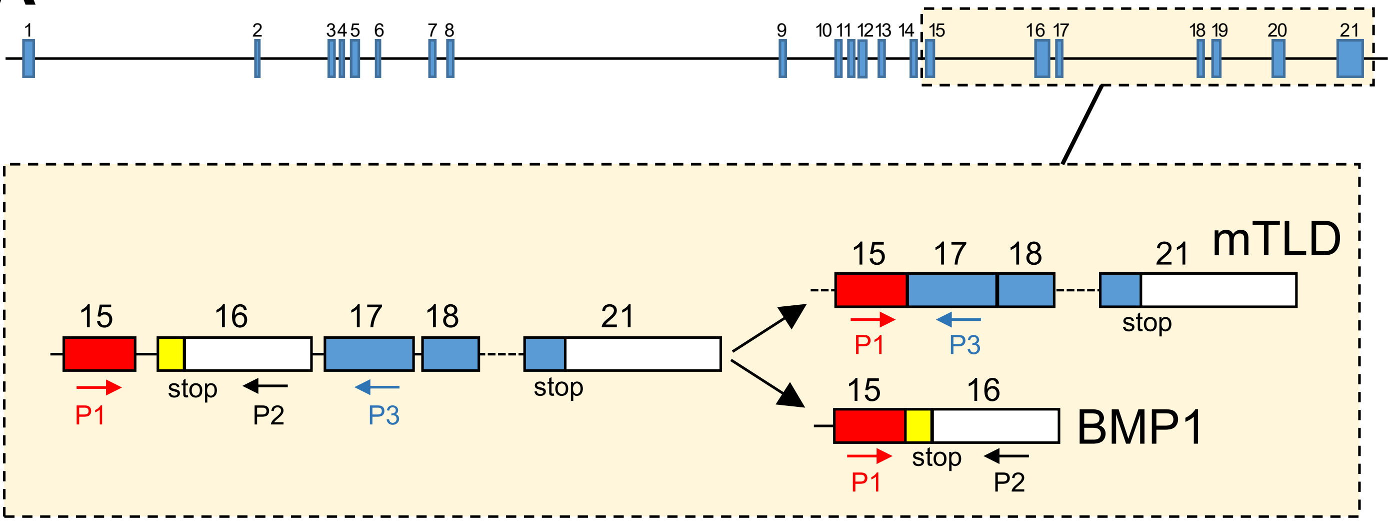


**B**

| Primer name | Sequence |
|---|---|
| Primer exon 15 forward (P1) | TCTGTGGTTCTGAGAAGCCC |
| Primer exon 16 reverse (P2) | CTCGGAATTTGAGCTGGTGG |
| Primer exon 17 reverse (P3) | TAACTGCCGAACGTGTTGAC |

**C**

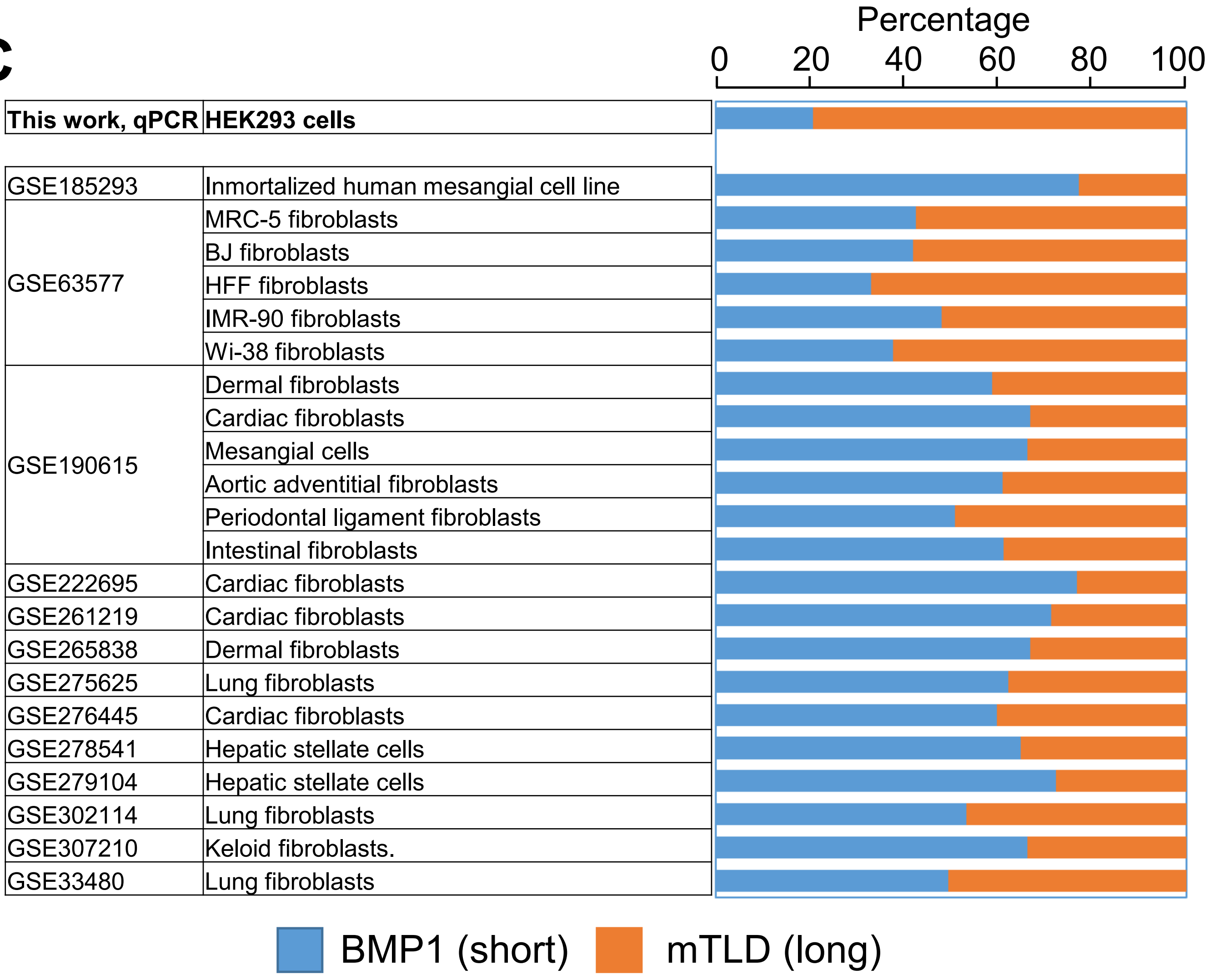


**Supplementary Figure 3**

### Supplementary Figure 5

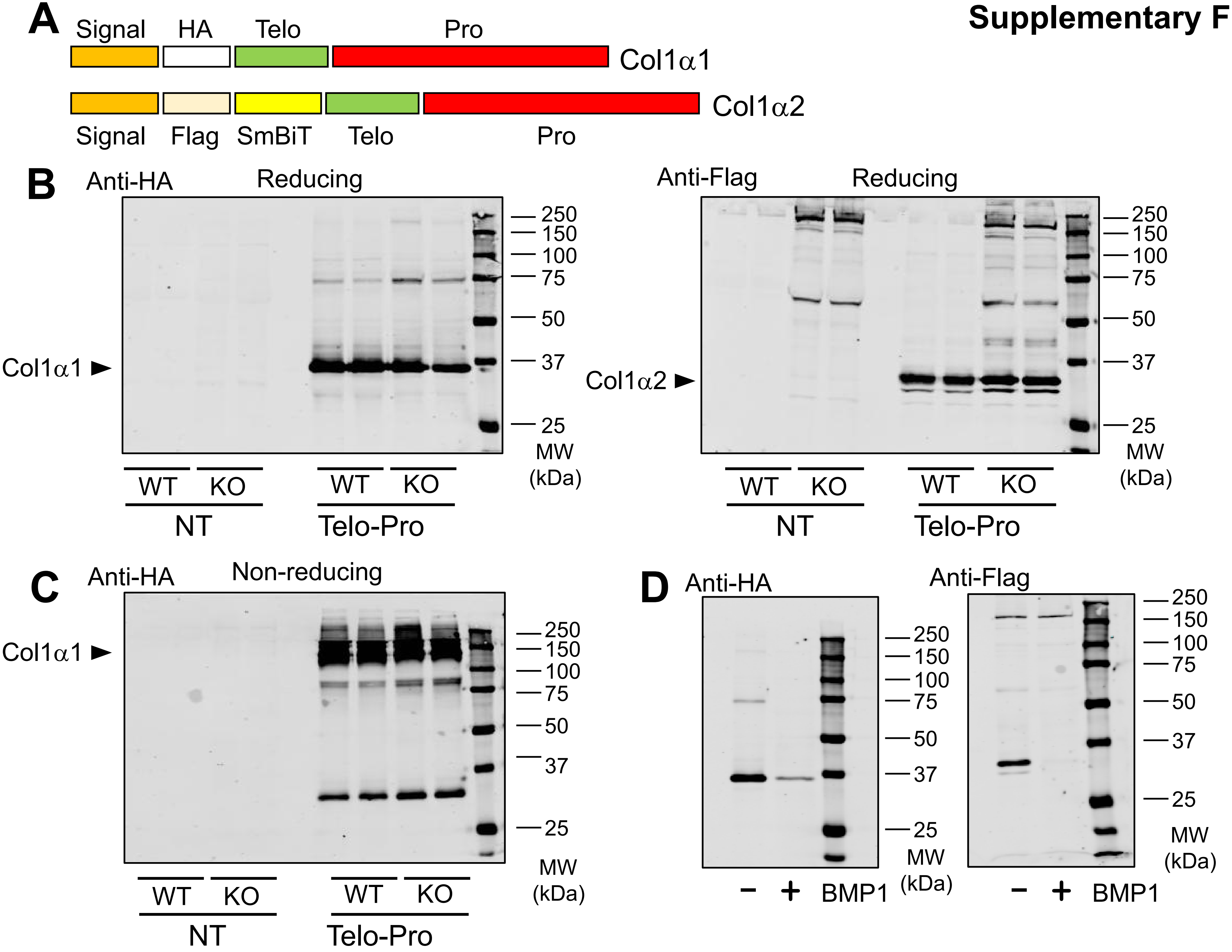

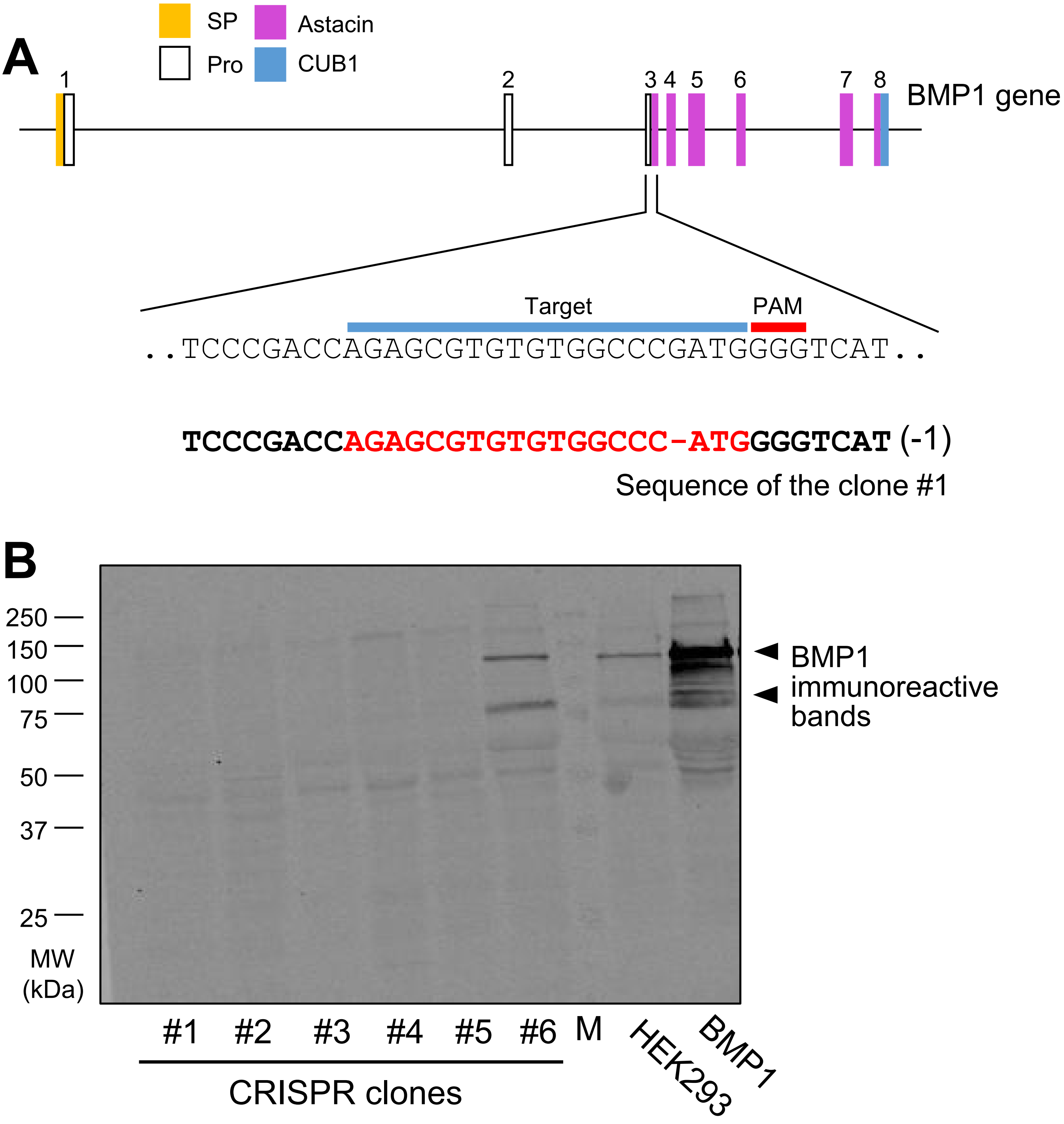


**Supplementary Figure 4**

| GEO Project ID | SRA project | SRA sample ID | Description | Tissue | % BMP1 (short) | % mTLD (long) | Mean short | Mean long |
|---|---|---|---|---|---|---|---|---|
| GSE185293 | SRX12485544 | SRR16200958 | Inmortalized human mesangial cell line. | kidney | 73.90 | 26.10 | 77.17 | 22.83 |
| | SRX12485546 | SRR16200959 | | | 75.90 | 24.10 | | |
| | SRX12485548 | SRR16200960 | | | 81.70 | 18.30 | | |
| GSE63577 | SRX766241 | SRR1660534 | MRC-5 fibroblasts | lung | 46.00 | 54.00 | 42.63 | 57.37 |
| | SRX766242 | SRR1660535 | | | 37.50 | 62.50 | | |
| | SRX766243 | SRR1660536 | | | 44.40 | 55.60 | | |
| | SRX766247 | SRR1660540 | BJ fibroblasts | skin | 39.10 | 60.90 | 42.03 | 57.97 |
| | SRX766248 | SRR1660541 | | | 50.00 | 50.00 | | |
| | SRX766249 | SRR1660542 | | | 37.00 | 63.00 | | |
| | SRX766253 | SRR1660546 | HFF fibroblasts | lung | 29.10 | 70.90 | 33.03 | 66.97 |
| | SRX766254 | SRR1660547 | | | 36.70 | 63.30 | | |
| | SRX766255 | SRR1660548 | | | 33.30 | 66.70 | | |
| | SRX766259 | SRR1660552 | IMR-90 fibroblasts | lung | 51.70 | 48.30 | 47.97 | 52.03 |
| | SRX766260 | SRR1660553 | | | 46.70 | 53.30 | | |
| | SRX766261 | SRR1660554 | | | 45.50 | 54.50 | | |
| | SRX766265 | SRR1660558 | Wi-38 fibroblasts | lung | 37.00 | 63.00 | 37.67 | 62.33 |
| | SRX766266 | SRR1660559 | | | 41.70 | 58.30 | | |
| | SRX766267 | SRR1660560 | | | 34.30 | 65.70 | | |
| | SRX13372265 | SRR17190440 | Dermal fibroblasts | Skin | 47.20 | 52.80 | 58.79 | 41.21 |
| | SRX13372266 | SRR17190441 | | | 62.50 | 37.50 | | |
| | SRX13372267 | SRR17190442 | | | 62.30 | 37.70 | | |
| | SRX13372268 | SRR17190443 | | | 53.20 | 46.80 | | |
| | SRX13372269 | SRR17190444 | | | 50.00 | 50.00 | | |
| | SRX13372270 | SRR17190445 | | | 61.50 | 38.50 | | |
| | SRX13372271 | SRR17190446 | | | 61.90 | 38.10 | | |
| | SRX13372272 | SRR17190447 | | | 71.70 | 28.30 | | |
| | SRX13372281 | SRR17190456 | Cardiac fibroblasts | Heart | 78.90 | 21.10 | 66.93 | 33.08 |
| | SRX13372282 | SRR17190457 | | | 60.00 | 40.00 | | |
| | SRX13372283 | SRR17190458 | | | 64.40 | 35.60 | | |
| | SRX13372284 | SRR17190459 | | | 84.90 | 15.10 | | |
| | SRX13372285 | SRR17190460 | | | 68.30 | 31.70 | | |
| | SRX13372286 | SRR17190461 | | | 60.30 | 39.70 | | |
| | SRX13372287 | SRR17190462 | | | 50.00 | 50.00 | | |
| | SRX13372288 | SRR17190463 | | | 68.60 | 31.40 | | |
| | SRX13372321 | SRR17190496 | Mesangial cells | Kidney | 56.20 | 43.80 | 66.26 | 33.74 |
| | SRX13372322 | SRR17190497 | | | 57.10 | 42.90 | | |
| | SRX13372323 | SRR17190498 | | | 59.40 | 40.60 | | |
| | SRX13372324 | SRR17190499 | | | 75.00 | 25.00 | | |
| | SRX13372325 | SRR17190500 | | | 74.40 | 25.60 | | |
| | SRX13372326 | SRR17190501 | | | 69.20 | 30.80 | | |

| | | | | | | | | |
|---|---|---|---|---|---|---|---|---|
| GSE190615 | SRX13372327 | SRR17190502 | | | 69.20 | 30.80 | | |
| | SRX13372328 | SRR17190503 | | | 69.60 | 30.40 | | |
| | SRX13372369 | SRR17190544 | Aortic adventitial fibroblasts | Heart | 66.70 | 33.30 | 61.06 | 38.94 |
| | SRX13372370 | SRR17190545 | | | 66.00 | 34.00 | | |
| | SRX13372371 | SRR17190546 | | | 60.40 | 39.60 | | |
| | SRX13372372 | SRR17190547 | | | 54.40 | 45.60 | | |
| | SRX13372373 | SRR17190548 | | | 61.70 | 38.30 | | |
| | SRX13372374 | SRR17190549 | | | 67.30 | 32.70 | | |
| | SRX13372375 | SRR17190550 | | | 50.00 | 50.00 | | |
| | SRX13372376 | SRR17190551 | | | 62.00 | 38.00 | | |
| | SRX13372377 | SRR17190552 | Periodontal ligament fibroblasts | Bone | 56.20 | 43.80 | 50.94 | 49.06 |
| | SRX13372378 | SRR17190553 | | | 63.60 | 36.40 | | |
| | SRX13372379 | SRR17190554 | | | 47.40 | 52.60 | | |
| | SRX13372380 | SRR17190555 | | | 50.00 | 50.00 | | |
| | SRX13372381 | SRR17190556 | | | 62.50 | 37.50 | | |
| | SRX13372382 | SRR17190557 | | | 63.20 | 36.80 | | |
| | SRX13372383 | SRR17190558 | | | 30.00 | 70.00 | | |
| | SRX13372384 | SRR17190559 | | | 34.60 | 65.40 | | |
| | SRX13372417 | SRR17190592 | Intestinal fibroblasts | Intestine | 45.00 | 55.00 | 61.25 | 38.75 |
| | SRX13372418 | SRR17190593 | | | 70.70 | 29.30 | | |
| | SRX13372419 | SRR17190594 | | | 58.10 | 41.90 | | |
| | SRX13372420 | SRR17190595 | | | 70.80 | 29.20 | | |
| | SRX13372421 | SRR17190596 | | | 59.20 | 40.80 | | |
| | SRX13372422 | SRR17190597 | | | 67.80 | 32.20 | | |
| | SRX13372423 | SRR17190598 | | | 67.30 | 32.70 | | |
| | SRX13372424 | SRR17190599 | | | 51.10 | 48.90 | | |
| GSE222695 | SRX19008313 | SRR23054998 | Cardiac fibroblasts | Heart | 75.20 | 24.80 | | |
| | SRX19008312 | SRR23054999 | | | 78.30 | 21.70 | 76.75 | 23.25 |
| GSE261219 | SRX23885546 | SRR28276169 | Cardiac fibroblasts | Heart | 63.50 | 36.50 | | |
| | SRX23885545 | SRR28276170 | | | 73.70 | 26.30 | 71.30 | 28.70 |
| | SRX23885544 | SRR28276171 | | | 68.90 | 31.10 | | |
| GSE265838 | SRX24361820 | SRR28798044 | Dermal fibroblasts | Skin | 66.40 | 33.60 | | |
| | SRX24361819 | SRR28798045 | | | 70.00 | 30.00 | 66.80 | 33.20 |
| | SRX24361818 | SRR28798046 | | | 63.60 | 36.40 | | |
| GSE275625 | SRX25823965 | SRR30397693 | Lung fibroblasts | lung | 69.00 | 31.00 | | |
| | SRX25823964 | SRR30397694 | | | 55.30 | 44.70 | 62.15 | 37.85 |
| GSE276445 | SRX25977110 | SRR30553780 | Cardiac fibroblasts | Heart | 61.80 | 38.20 | | |
| | SRX25977107 | SRR30553783 | | | 58.60 | 41.40 | | |
| | SRX25977104 | SRR30553786 | | | 64.00 | 36.00 | 59.77 | 40.23 |
| | SRX25977101 | SRR30553789 | | | 52.70 | 47.30 | | |
| | SRX25977098 | SRR30553792 | | | 61.70 | 38.30 | | |

| | | | | | | | | |
|---|---|---|---|---|---|---|---|---|
| | SRX25977095 | SRR30553795 | | | 59.80 | 40.20 | | |
| GSE278541 | SRX26253375 | SRR30854995 | Hepatic stellate cells | Liver | 74.60 | 25.40 | | |
| | SRX26253367 | SRR30855003 | | | 55.10 | 44.90 | 64.85 | 35.15 |
| GSE279104 | SRX26323944 | SRR30920939 | Hepatic stellate cells | Liver | 68.30 | 31.70 | 72.33 | 27.68 |
| | SRX26323943 | SRR30920940 | | | 79.50 | 20.50 | | |
| | SRX26323942 | SRR30920941 | | | 70.10 | 29.90 | | |
| | SRX26323941 | SRR30920942 | | | 80.50 | 19.50 | | |
| | SRX26323937 | SRR30920946 | | | 66.20 | 33.80 | | |
| | SRX26323928 | SRR30920955 | | | 74.50 | 25.50 | | |
| | SRX26323927 | SRR30920956 | | | 74.80 | 25.20 | | |
| | SRX26323926 | SRR30920957 | | | 80.20 | 19.80 | | |
| | SRX26323925 | SRR30920958 | | | 68.30 | 31.70 | | |
| | SRX26323924 | SRR30920959 | | | 81.60 | 18.40 | | |
| | SRX26323912 | SRR30920971 | | | 68.10 | 31.90 | | |
| | SRX26323911 | SRR30920972 | | | 81.00 | 19.00 | | |
| | SRX26323910 | SRR30920973 | | | 79.80 | 20.20 | | |
| | SRX26323909 | SRR30920974 | | | 76.70 | 23.30 | | |
| | SRX26323908 | SRR30920975 | | | 65.00 | 35.00 | | |
| | SRX26323907 | SRR30920976 | | | 79.10 | 20.90 | | |
| | SRX26323893 | SRR30920990 | | | 75.00 | 25.00 | | |
| | SRX26323892 | SRR30920991 | | | 68.00 | 32.00 | | |
| | SRX26323890 | SRR30920993 | | | 52.70 | 47.30 | | |
| | SRX26323889 | SRR30920994 | | | 62.10 | 37.90 | | |
| | SRX26323888 | SRR30920995 | | | 71.00 | 29.00 | | |
| | SRX26323887 | SRR30920996 | | | 67.10 | 32.90 | | |
| | SRX26323886 | SRR30920997 | | | 65.40 | 34.60 | | |
| | SRX26323885 | SRR30920998 | | | 80.80 | 19.20 | | |
| GSE302114 | SRX29599265 | SRR34438170 | Lung fibroblasts | Lung | 57.80 | 42.20 | 53.33 | 46.67 |
| | SRX29599264 | SRR34438171 | | | 59.50 | 40.50 | | |
| | SRX29599263 | SRR34438172 | | | 42.70 | 57.30 | | |
| GSE307210 | SRX30350002 | SRR35247104 | Keloid fibroblasts. | Skin | 65.00 | 35.00 | 66.30 | 33.70 |
| | SRX30350001 | SRR35247105 | | | 66.20 | 33.80 | | |
| | SRX30350000 | SRR35247106 | | | 67.70 | 32.30 | | |
| GSE33480 | SRX159832 | SRR521527 | Lung fibroblasts | lung | 54.60 | 45.40 | 49.53 | 50.48 |
| | SRX159832 | SRR521528 | | | 40.30 | 59.70 | | |
| | SRX159832 | SRR521529 | | | 48.70 | 51.30 | | |
| | SRX159832 | SRR521530 | | | 54.50 | 45.50 | | |
